# Inferring state-dependent diversification rates using approximate Bayesian computation (ABC)

**DOI:** 10.1101/2023.10.14.562317

**Authors:** Shu Xie, Luis Valente, Rampal S. Etienne

**Author notes:** CORRESPONDING AUTHOR *Shu Xie Email:.

## Abstract

State-dependent speciation and extinction (SSE) models provide a framework for quantifying whether species traits have an impact on evolutionary rates and how this shapes the variation in species richness among clades in a phylogeny. However, SSE models are becoming increasingly complex, limiting the application of likelihood-based inference methods. Approximate Bayesian computation (ABC), a likelihood-free approach, is a potentially powerful alternative for estimating parameters. One of the key challenges in using ABC is the selection of efficient summary statistics, which can greatly affect the accuracy and precision of the parameter estimates. In state-dependent diversification models, summary statistics need to capture the complex relationships between rates of diversification and species traits. Here, we develop an ABC framework to estimate state-dependent speciation, extinction and transition rates in the BiSSE (binary state dependent speciation and extinction) model. Using different sets of candidate summary statistics, we then compare the inference ability of ABC with that of using likelihood-based maximum likelihood (ML) and Markov chain Monte Carlo (MCMC) methods. Our results show the ABC algorithm can accurately estimate state-dependent diversification rates for most of the model parameter sets we explored. The inference error of the parameters associated with the species-poor state is larger with ABC than in the likelihood estimations only when the speciation rate is highly asymmetric between the two states (*λ*_1_ / *λ*_0_ = 5). Furthermore, we find that the combination of normalized lineage-through-time (nLTT) statistics and phylogenetic signal in binary traits (Fitz and Purvis’s *D*) constitute efficient summary statistics for the ABC method. By providing insights into the selection of suitable summary statistics, our work aims to contribute to the use of the ABC approach in the development of complex state-dependent diversification models, for which a likelihood is not available.

## Introduction

Detecting the factors that underlie variations in diversification rate is a major topic of research in evolutionary biology, as it may reveal the causes of unevenness of species richness among clades and geographical regions. Numerous studies highlight the important role of traits in shaping speciation and extinction rates. For example, the presence of the nectar spur in flowering plants (angiosperms), which facilitates pollination and reproductive success, is associated with rapid diversification in the clades of spurred species (Armbruster, 2014; Fernández-Mazuecos et al., 2019). Traits such as this are hypothesized to affect evolutionary processes by determining the interactions between species, as well as how species respond to the environment (Wiens, 2017; Li & Wiens, 2022).

Over the past years, phylogenetic comparative methods to test evolutionary hypotheses have been developed rapidly (Miles & Dunham, 1993; Adams, 2013). A popular class of phylogenetic tools are state-dependent speciation and extinction (SSE) models, which aim to investigate how species traits shape variation in diversification and richness, and to infer the rates of evolutionary processes (speciation, extinction and transitions between character states) (Maddison et al., 2007). BiSSE (binary-state speciation and extinction) is the original SSE model, and has been expanded to consider quantitative (QuaSSE, Fitzjohn, 2010), geographic (GeoSSE, Goldberg et al., 2011), and multiple categorical states (MuSSE, Fitzjohn, 2012). However, these models have been shown to suffer from a high risk of false positives, which are likely to attribute differential diversification rate to trait-dependence. In order to reduce this type I error in the SSE models, Beaulieu & O’Meara (2016) introduced the HiSSE (hidden-state-dependent speciation and extinction) model incorporating the effect of hidden states, which further promotes the subsequent development of state-dependent diversification models. More recently, Herrera-Alsina et al., (2019) introduced the SecSSE (several examined and concealed states-dependent speciation and extinction) framework accounting for multiple traits (observed and hidden) and multiple trait states. These models have been applied in numerous empirical studies to identify the correlation between trait states and diversification rates (Onstein et al., 2017; Pyron & Burbrink, 2012; Rolland, Condamine, et al., 2014).

The most commonly used methods for estimating parameters in the current SSE models are likelihood-based inference approaches, such as maximum likelihood or Bayesian inference. In general, the likelihood calculation of these models relies on a set of ordinary differential equations (ODE) for two core sets of probabilities: 1) the probabilities of observing the phylogeny and associated character states at tips evolved from a lineage in each possible trait state at a time in the past, and 2) the extinction probabilities, i.e. the probabilities of a lineage in each state at a time in the past having no extant descendants at present (Maddison et al., 2007; FitzJohn, 2012). In recent years, researchers have explored the power of parameter estimations of existing models (Davis et al., 2013; Rabosky & Goldberg, 2015), and strived to improve the accuracy of the likelihood computing approaches (Louca & Pennell, 2020; Laudanno et al., 2021; Vasconcelos et al., 2022). However, SSE models have become increasingly complex, from considering a single trait with binary states (BiSSE) to multiple traits with hidden states (SecSSE). The computational cost and intractability limit the application of likelihood-based inference methods.

Approximate Bayesian computation (ABC) is a powerful alternative for estimating parameters when the likelihood is difficult to compute (Csilléry et al., 2010; Beaumont, 2019). ABC is a simulation-based Bayesian approach to find the parameters that can generate data close to the target (observed data) by evaluating the similarity of a set of summary statistics between simulated and observed data (Tavare et al., 1997). Numerous methods have been developed to address the challenges of improving the efficiency of the ABC estimation. A series of efficient ABC algorithms have been expanded based on the simple rejection algorithm, such as incorporating Markov chain Monte Carlo (MCMC) (Marjoram et al., 2003), population Monte Carlo (PMC) (Beaumont et al., 2009) and sequential Monte Carlo (SMC) (Toni et al., 2009). The ABC-MCMC algorithm takes advantage of the Markov chain Monte Carlo techniques to explore the parameter space by sampling from Markov Chains, where the proposal distribution remains static throughout the sampling processes. ABC-PMC uses a population of particles that are iteratively updated to approximate posterior distribution. However, it may be challenging in ABC-MCMC and ABC-PMC to achieve convergence and explore high-dimensional parameter space. ABC-SMC combines the strengths of these two methods and performs better in such cases. The algorithm systematically improves the approximation of the posterior distribution through a series of intermediate distributions, which allows to converge to the posterior faster reducing the usage of computational resources and offering improved parameter estimation in complex models.

The developments of ABC methods have facilitated the application of ABC approaches in a broad field of studies. However, while these methods have been widely used in different fields, including ecological and evolutionary studies (Beaumont, 2010), the applications in trait evolution or diversification analysis are still very limited. Existing trait-related studies using ABC methods focus on detecting the impact of environment or species interactions on trait evolution or mapping trait evolution on given phylogenies (Janzen et al., 2016; Bartoszek & Lio, 2019; Xu et al., 2021), but no study has yet tried to apply ABC approaches to study the effect of trait dynamics on diversification rates.

ABC methods are sensitive to the selection of the summary statistics, therefore, it is necessary to find powerful summary statistics that cover the maximum amount of the information in the data (Sirén & Kaski, 2020). Efficient summary statistics for SSE models should ideally capture information on two aspects: the shape of the phylogenetic tree and the distribution of traits. Currently, few statistics have been synthesized to describe the dynamics of binary or multi-state categorical traits along phylogenies. Pagel’s *λ* (1999) and Blomberg’s *K* (2003) are the classic measures of phylogenetic signal, which evaluates how phylogenetic distance shapes the distribution of traits among species, as well as the frequency of trait changes along a phylogeny. However, these two statistics are designed for continuous trait data, and cannot be calculated for discrete trait data because it is not possible to calculate variances and co-variances from the trait distribution (Borges et al., 2019). More recently, two statistics (*D* and *δ*) were derived to measure phylogenetic signal particularly for binary or categorical traits (Fritz & Purvis, 2010; Borges et al., 2019). In addition, some efficient statistics (e.g., normalized Lineage Through Time (nLTT)) employed in phylogenetic ABC analyses of diversification (Janzen et al., 2015), as well as statistics applied to measure the phylogenetic diversity (mean pairwise distance (MPD), mean nearest taxon distance (MNTD)), can also be extended to measure the distribution of a trait along phylogenies by separately analyzing species with the same trait state. However, there is yet to be an evaluation of the performance of those summary statistics in trait state-dependent models.

Due to the high efficiency and accuracy in estimation, here, we use the ABC-SMC framework to estimate parameters in the simplest state-dependent model BiSSE, and test the inference performance by comparing the inference error of the parameters using the ABC-SMC algorithm and two likelihood-based approaches: maximum likelihood estimation (MLE) and Markov chain Monte Carlo (MCMC). Finally, we investigate the performance of a set of phylogenetic and trait related summary statistics to select the most efficient combination for estimating evolutionary rates of state-dependent models.

## Methods

### Trait-dependent simulation

The BiSSE (binary state speciation and extinction) model simulates diversification dynamics according to a birth-death model where traits shape diversification rates. The model considers an evolving binary trait with two states, 0 and 1 (e.g., presence or absence of a specific trait). It assumes that the per lineage rates of speciation (*λ_0_*, *λ_1_*) and per lineage extinction (*μ_0_*, *μ_1_*) depends on the trait state of a species. It also allows transitions between states (*q_01_*, *q_10_*). Based on this model, we simulated “observed” phylogenetic trees and trait states at tips, under a series of parameter scenarios considering symmetric (equal rates between trait states) or asymmetric (rates differ between trait states), speciation, extinction and transition rates (Table 1). To simplify the analysis, we set only one of the three pairs of rates to be asymmetric in each scenario. For each scenario we simulated 50 replicates, and in total we produced 350 phylogenetic trees with trait data as observed data for parameter inference. To avoid extremely large trees, we set a constraint with a maximum of 500 species in total, and for trait dependence to make sense (i.e., we would not expect to do an SSE-analysis when only one state is observed), we imposed the constraint that at least one species is present for each state at the end of the simulation.

**Table 1.**
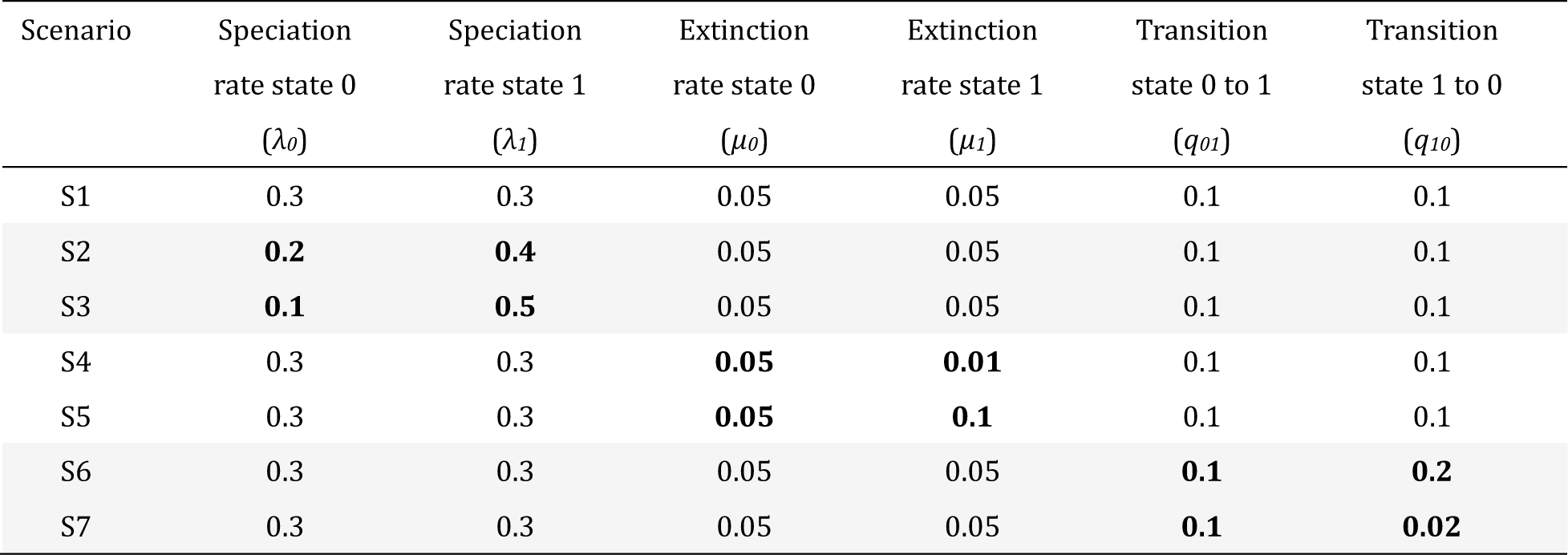
Parameter sets used to generate the observed data via simulations of the BiSSE model. In total, seven parameter combinations were used (seven scenarios), including a symmetric scenario as control (scenario 1), and two asymmetric scenarios for each pair of rates.

| Scenario | Speciation<br>rate state 0<br>($\lambda_0$) | Speciation<br>rate state 1<br>($\lambda_1$) | Extinction<br>rate state 0<br>($\mu_0$) | Extinction<br>rate state 1<br>($\mu_1$) | Transition<br>state 0 to 1<br>($q_{01}$) | Transition<br>state 1 to 0<br>($q_{10}$) |
| --- | --- | --- | --- | --- | --- | --- |
| S1 | 0.3 | 0.3 | 0.05 | 0.05 | 0.1 | 0.1 |
| S2 | **0.2** | **0.4** | 0.05 | 0.05 | 0.1 | 0.1 |
| S3 | **0.1** | **0.5** | 0.05 | 0.05 | 0.1 | 0.1 |
| S4 | 0.3 | 0.3 | **0.05** | **0.01** | 0.1 | 0.1 |
| S5 | 0.3 | 0.3 | **0.05** | **0.1** | 0.1 | 0.1 |
| S6 | 0.3 | 0.3 | 0.05 | 0.05 | **0.1** | **0.2** |
| S7 | 0.3 | 0.3 | 0.05 | 0.05 | **0.1** | **0.02** |

### Likelihood-based estimation through likelihood maximization and Bayesian MCMC

The BiSSE model allows likelihood calculation (Maddison et al., 2007), where the likelihood indicates the probability of observing the binary trait data on the phylogenetic tree with given parameters. The likelihood is calculated based on a set of ordinary differential equations (Maddison et al., 2007):

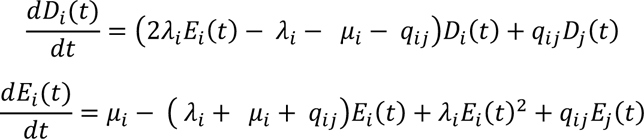

where *D_i_(t)* describes the probability of observing the phylogeny and associated character states at present evolved from a lineage in a particular trait state *i* at time *t*, and *E_i_*(*t*) describes the probability of a lineage that has no descendants at present, and *λ_i_, μ_i_, q_ij_* represent state-dependent speciation, extinction and character transition rates respectively (Maddison et al., 2007; FitzJohn, 2012).

Parameters can then be estimated using maximum likelihood estimation (MLE), which yields only a point estimate. To compare the ABC results with the likelihood-based approach, we developed a full (i.e., using the likelihood) Bayesian analysis using Markov chain Monte Carlo (MCMC), under the same assumptions of prior distributions as the ABC (see below). To obtain a stable and convergent MCMC chain, we ran 1,000,000 iterations after 100,000 iterations of burn-in. For each data set, we estimated all six parameters regardless of the symmetry of the generating rates. The simulations and likelihood calculations were performed using the R package *secsse* (Herrera-Alsina et al., 2023). The reason for using the *secsse* package is that the SecSSE model reduces exactly to BiSSE when there are no hidden states and the examined states are binary, but the range of application is much broader than BiSSE. As the combination of HiSSE and MuSSE, the SecSSE model takes into account hidden states, which improves the accuracy of detecting trait dependencies in diversification rates, as well as breaking the constraints on the number of traits and trait states in preceding SSE models. Therefore, it can accurately generate BiSSE simulations and estimates, and facilitates further testing in the more complex conditions.

### ABC-SMC estimation

We performed a sequential Monte Carlo algorithm (ABC-SMC) to estimate parameters for each observed data. The algorithm we used was derived from the original ABC-SMC algorithm introduced by Toni et al. (2009) (Box 1). For the ABC algorithm, we used uniform prior distributions U (0,1). The algorithm starts by sampling a series of parameter sets (particles) from the prior distribution, and then simulates datasets with these parameters and computes the difference in summary statistic between the observed data and the simulated data. This was repeated until this difference was smaller than a threshold ɛ. We used an iteratively adaptive method choosing ever-decreasing thresholds for each iteration, by specifying the median values of the summary statistic distance from the previous iteration, which means the decreasing thresholds depend on the position of the accepted particles in the previous iteration. This is more efficient than using a given linearly or exponentially decreasing pattern. We used 500 particles per iteration and the algorithm was assumed to generate a converged posterior at an acceptance rate of 1 in 500. The ABC and MCMC algorithms used in this study were implemented in the R package DivABC, which is available on Github (github.com/xieshu95/DivABC).

#### Box 1. The ABC SMC algorithm

S1: Initialize thresholds *ɛ_1_*
S2: Set iteration *t* = 1
  For particle *i* = 1,…, *N*
    Repeat
      Sample *θ_1_^i^* from the prior *π*(*θ*).
      Simulate phylogeny *P_1_^i^* ∼ *θ_1_^i^*.
      Calculate distance *D_1_^i^* (*SS*) = |*SS*(*P_1_^i^*) – *SS*(*P*_0_)|.
    until *D_1_^i^* (*SS*) < *ɛ_1_*
      Calculate weight: 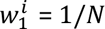
  Calculate threshold for next iteration: *ɛ_2_* = median {*D_1_^1^* (*SS*), *D_1_^2^* (*SS*)… *D_1_^N^* (*SS*)}
S3: *t* = 2,…,*T*
  For particle *i* = 1,…, *N*
    Repeat
      Sample *θ_t_^i^* from population {*θ_t_*_-1_} with weight 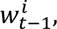 and perturb *θ_t_*^i^ ∼*N*(0, 0.01).
      Simulate phylogeny *P_t_^i^* ∼ *θ_t_^i^*.
      Calculate distance *D_t_^i^* (*SS*) = |*SS*(*P_t_^i^*) – *SS*(*P*_0_)|.
    until *D_t_^i^* (*SS*) < *ɛ_t_*
      Add *θ_t_^i^* to the population {*θ_t_*}
      Calculate weight: 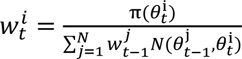
  Normalize the weights.
  Calculate threshold for next iteration: *ɛ_t_* _+ 1_ = median {*D_t_^1^* (*SS*), *D_t_^2^* (*SS*)… *D_t_^N^* (*SS*)}

where *N* is the total number of the particles needed for generation *t*, and *T* is the total number of iterations before the algorithm stop. *ɛ_1_* …*ɛ_T_* means the sequence of decreasing tolerance threshold from iteration 1 to T. *θ_t_*^i^ means the particle *i* of iteration *t*, and 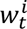 is the weight of this particle. *P_t_^i^* is the simulated data with the particle *θ_t_*^i^, and *SS*(*P_t_^i^*) is the calculated summary statistic of the simulation.

### Summary statistics

We tested the usefulness and efficiency of a set of summary statistics for improving ABC performance. In total, we used six summary statistics to describe the phylogenetic dynamics and trait evolution along the phylogeny, which are: 1) NLTT (normalized lineage-through-time), 2) MPD (mean pairwise distance), 3) MNTD (mean nearest taxon distance), 4) Colless index, 5) tip ratio, 6) phylogenetic signal *D*. NLTT is a summary statistic known to be efficient in phylogenetic analyses of diversification, which has been shown a better performance than classic statistics (e.g., phylogenetic diversity (PD)) in different types of birth-death models (Janzen et al., 2015). The original nLTT statistic does not capture trait information, so we developed a method to calculate the nLTT statistic of each trait state by trimming branches from the phylogenetic tree with the other trait state at the tip. We note that the combination of the two resulting trimmed trees may not be exactly equivalent to the original tree (Fig. 1). We calculated the nLTT statistics for the entire tree (nLTT_total_), for the trimmed tree with only state 0 (nLTT_0_), and the trimmed tree with only state 1 (nLTT_1_). As illustrated in Fig 1, for the same phylogenetic tree, different trait distributions at the tips can lead to differences in nLTT_0_ or nLTT_1_. MPD and MNTD are metrics that have been commonly used to measure phylogenetic diversity (Webb, 2000), and Colless index is a widely used statistic to measure the balance of phylogenetic trees (Colless, 1995). Furthermore, we calculated the state-specific MPD, MNTD and Colless index in the same way as calculating nLTT_0_ or nLTT_1_ based on the trimmed tree with a single state. The tip ratio between binary states was calculated as the number of species with species-rich state divided by the number of species with species-poor state:

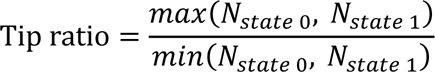

**Fig 1.**
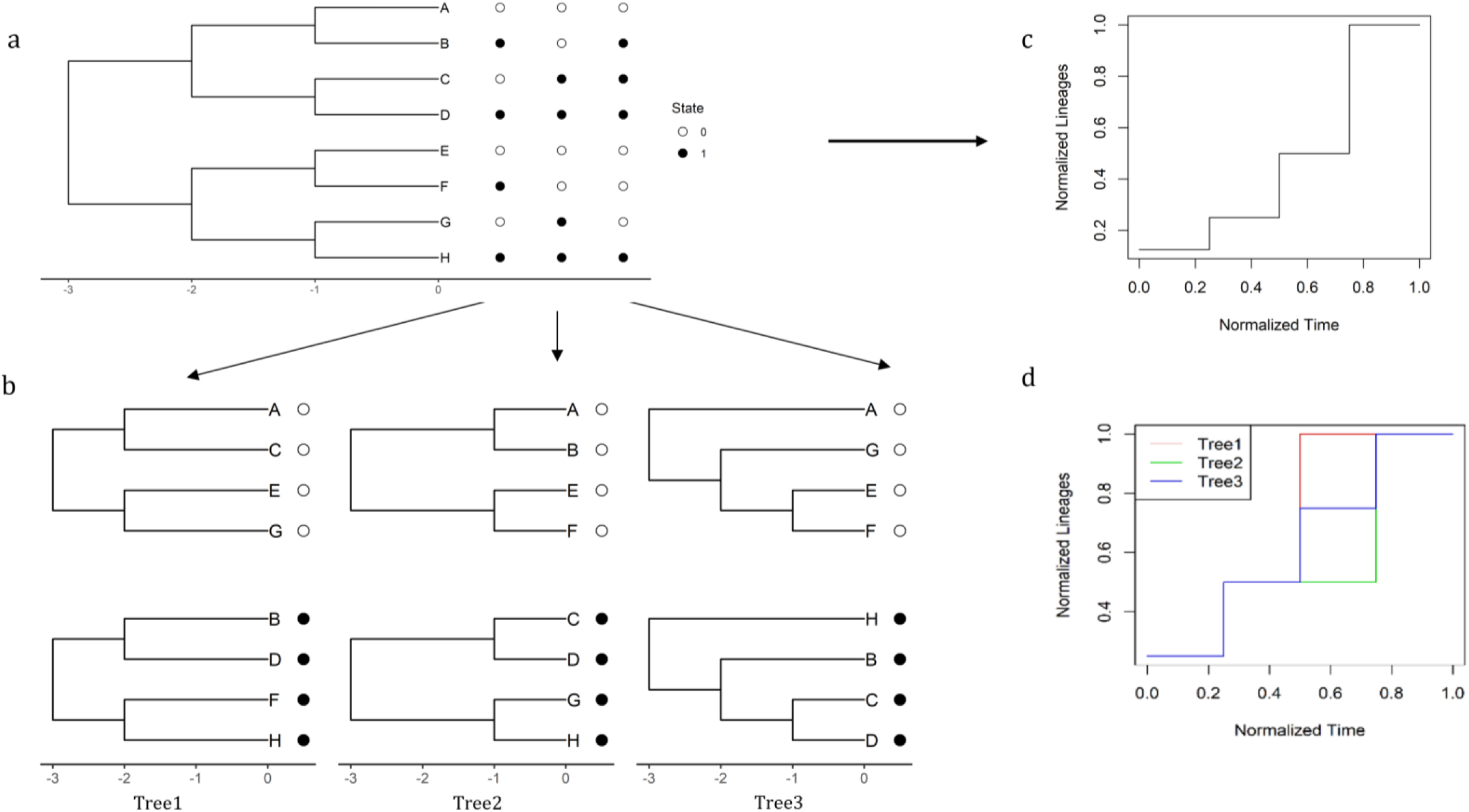
Hypothetical example of transforming an entire phylogeny with binary tip states into two trimmed trees with a single state by clustering the tips with the same states. a) example of a (balanced) tree with eight extant species at the present time, and three potential trait distribution at tips (other trait distributions are possible, but we show only three for the example). Blank circles represent state 0, and filled black circles represent state 1. b) shows the two reduced trees for each state depending on the trait distribution at the tips a). c) the plot of the nLTT for the entire phylogenetic tree in a). d) the plot of the nLTT for the trimmed trees with a single state in b), and here we show only the nLTT plot for one of the states, because the plot for the other state is equivalent in the example.

Another summary statistic we examined is *D* (Fritz & Purvis, 2010), which measures the phylogenetic signal of a given phylogeny with binary trait states. To calculate *D,* the total state difference between each sister clade along the phylogeny (∑*d_obs_*) (Fig 2) is computed, and is then scaled by the sum of state differences based on two permutations of the trait values. One is a random permutation that shuffles the tip state values (0 or 1) along the tree, generating a series of sums of state differences ∑*d_r_*, and the other simulates continuous trait evolution under the Brownian motion model, and discretizes the tip states into a binary trait, generating a series of sums of state difference ∑*d_b_*. Permutations are performed 1000 times for each tree. The statistic *D* is calculated as:

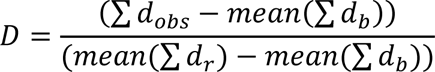

**Fig 2.**
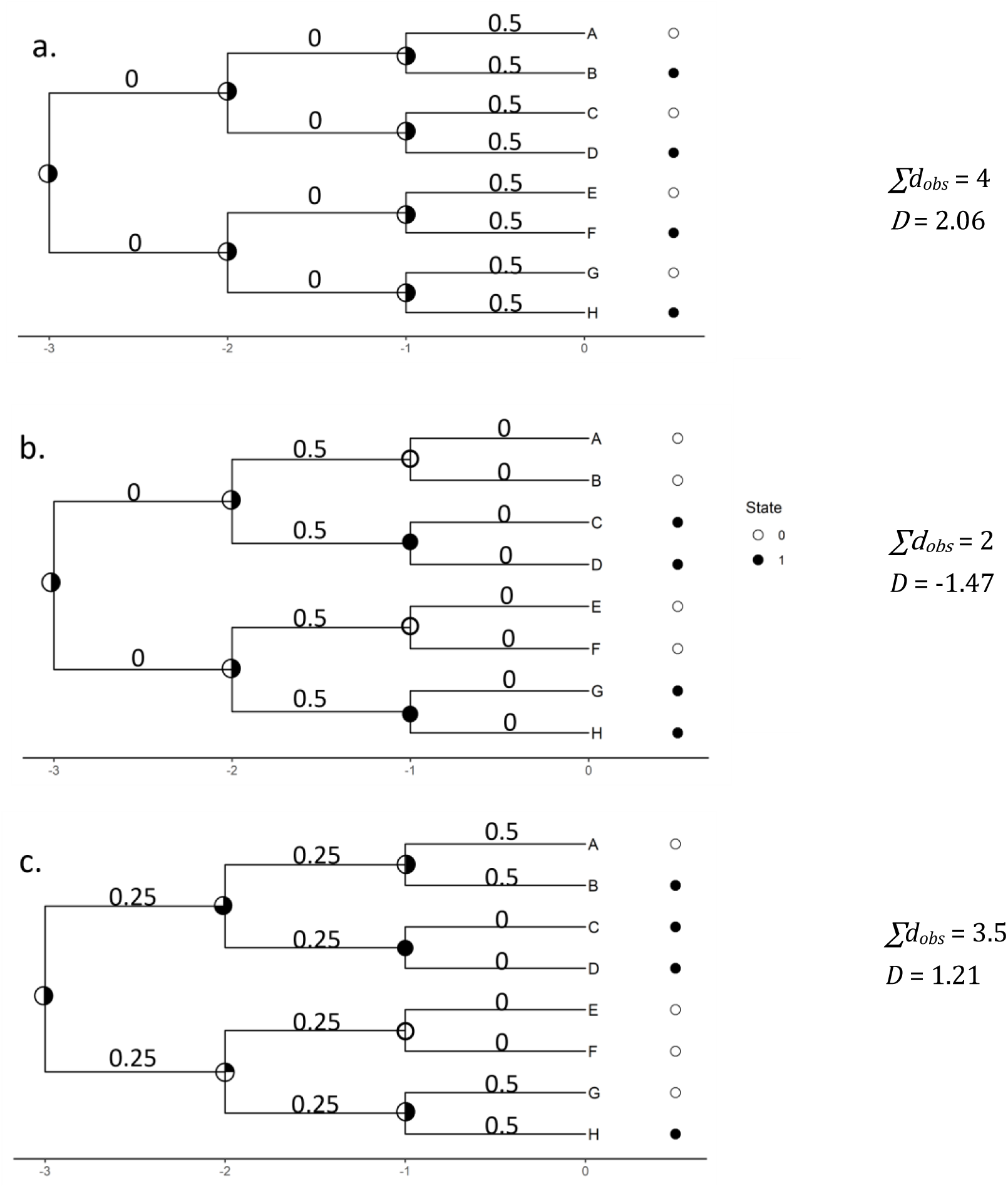
Illustration of calculating *∑d_obs_* and *D* under different phylogenetic patterns. A) is a phylogenetically overdispersed tree with binary sates evenly distributed at tips. B) and c) are two phylogenetic clumped trees. The circles at the tips indicate the observed trait states, and the circles at nodes indicate the probability of each ancestral state. To calculate *D*, the mean (∑d_r_) and mean (∑d_b_) are determined from 1000 permutations for each tree.

The ∑*d_obs_* is sensitive to the pattern of how traits evolve through the phylogeny (phylogenetically clumped or dispersed), and the *D* statistic can distinguish trait dynamics even under the same phylogeny (Fig 2).

To simplify the analysis, we did not test all the permutations among these six metrics, but manually selected the combinations of most interest (Table 2). We used the nLTT_total_ as the main measurement of the phylogenetic dynamics, however, this metric is insufficient for inferring diversification variance between states because of the lack of trait information. Therefore, we added other trait-related statistics respectively based on nLTT_total_, and compared the performance among the combinations to filter the most powerful statistics. The calculations of nLTT, MPD, MNTD and Colless index are implemented in the R package *treestats*, which is available on Github (github.com/thijsjanzen/treestats).

**Table 2.** Selected summary statistic combinations and their abbreviations.

| Combination | Summary statistic | Abbreviation |
| --- | --- | --- |
| 1 | $nLTT_{total}$ | nLTT |
| 2 | $D$ | $D$ |
| 3 | $nLTT_{total} + D$ | nLTT-$D$ |
| 4 | $nLTT_{total} + nLTT_1 + nLTT_2$ | nLTTs |
| 5 | $nLTT_{total} + nLTT_1 + nLTT_2 + D$ | nLTTs-$D$ |
| 6 | $nLTT_{total} + MPD_1 + MPD_2$ | nLTT-MPD |
| 7 | $nLTT_{total} + MNTD_1 + MNTD_2$ | nLTT-MNTD |
| 8 | $nLTT_{total} + colless_1 + colless_2$ | nLTT-colless |
| 9 | $nLTT_{total} + \text{tip ratio}$ | nLTT-ratio |

## Results

We compared the inference error of the different inference methods by calculating the relative distance between the true (generating) values and the estimations using the ABC, MCMC and MLE approaches. In the main text, we use the median of the posterior distributions representing the estimations from the ABC and MCMC algorithms to compare with the point estimation of MLE. The comparisons of the full posterior distributions are given in the Supplementary Material (Fig S3). To analyze the effect of different levels of trait dependence on parameter estimation, we divided the seven scenarios into three groups according to the asymmetry of different rates, which are 1) asymmetry in speciation (scenarios S1, S2 and S3); 2) asymmetry in extinction (scenarios S1, S4 and S5); 3) asymmetry in transition (scenarios S1, S6 and S7).

Our general conclusion is that using only the nLTT statistics in the ABC approach can lead to accurate estimates of state-dependent speciation and extinction rates, but produces a large inference error in estimating transition rates between states. However, the inference accuracy can be significantly increased by adding the summary statistic *D*. In this case, the ABC algorithm performs well in estimating all six parameters of the BiSSE model for most of the scenarios we investigated, and is comparable with the likelihood-based approaches with minor differences. The ABC methods only lead to relatively larger inference errors when the speciation rates are highly asymmetric between states (*λ*_1_ / *λ*_0_ = 5), but in this case the error occurs only in the rates associated with the state with fewer species. We will now discuss our results in detail.

### Statistics of the observed data

We calculated the tree size (as measured by the total number of species on the observed phylogenetic tree), tip ratio (the ratio of the diversity between the species-rich state and the species-poor state), and the number of tips with each state across all the observed datasets (350 trees). The observed phylogenetic trees generated under the seven scenarios show different patterns. Overall, the mean and the standard deviation of the size of the full tree are similar among the scenarios (Table 2), because of the constraint of the maximum number (500) of the species when generating observed data. However, the tip ratio becomes larger with an increasing level of asymmetry in speciation and transition rates. In addition, as the asymmetry level in speciation increasing, more observed trees with ancestral state 0 (with lower speciation rate) has been selected (Table 2).

**Table 2.** Main properties of the observed datasets. For tree size, tip ratio, number of tips with state 0 and state 1, we show the mean (standard deviation) across the 50 observed datasets for each scenario (obtained via simulations using the parameters of each scenario). The last two columns show the number of replicates with each ancestral state respectively.

| Scenario | Tree size | Tip ratio | N<sub>State0</sub> | N<sub>State1</sub> | Ancestral state 0 | Ancestral state 1 |
| --- | --- | --- | --- | --- | --- | --- |
| S1 | 222 (98) | 1.29 (0.28) | 113 (52) | 110 (51) | 25 | 25 |
| S2 | 230 (111) | 2.39 (0.83) | 70 (33) | 161 (82) | 34 | 16 |
| S3 | 232 (127) | 4.55 (2.48) | 46 (27) | 186 (102) | 42 | 8 |
| S4 | 241 (120) | 1.40 (0.35) | 109 (57) | 131 (68) | 28 | 22 |
| S5 | 195 (92) | 1.39 (0.27) | 110 (56) | 86 (40) | 28 | 22 |
| S6 | 233 (114) | 2.19 (0.65) | 158 (78) | 76 (38) | 25 | 25 |
| S7 | 246 (106) | 5.72 (3.78) | 46 (30) | 200 (91) | 22 | 28 |

### ABC with different summary statistics

Using different groups of summary statistics shows a large difference in the performance of parameter estimation of ABC. The nLTT statistic alone can accurately estimate speciation and extinction rates (Figs 3, 4, and 5) except when the generating rates of speciation greatly vary between binary states (i.e., scenario S3) (Figs 3, 4, and 5), and leads to large bias in net diversification rate estimates in this case (Fig S1). However, the inference errors and the variance in transition rates and net diversification rates among replicates are always large (Figs 3, 4, and 5), due to the lack of trait dynamic information along phylogenetic trees. The combination of nLTTs and nLTT-MNTD improves the inference accuracy in estimating speciation rates when there is high asymmetry in speciation (Fig 3), while the other combinations including nLTT (i.e., nLTT-MPD, nLTT-colless, and nLTT-ratio) show no significant improvement in estimations over using nLTT alone (Figs 3, 4, 5 and S1).

**Fig 3.**
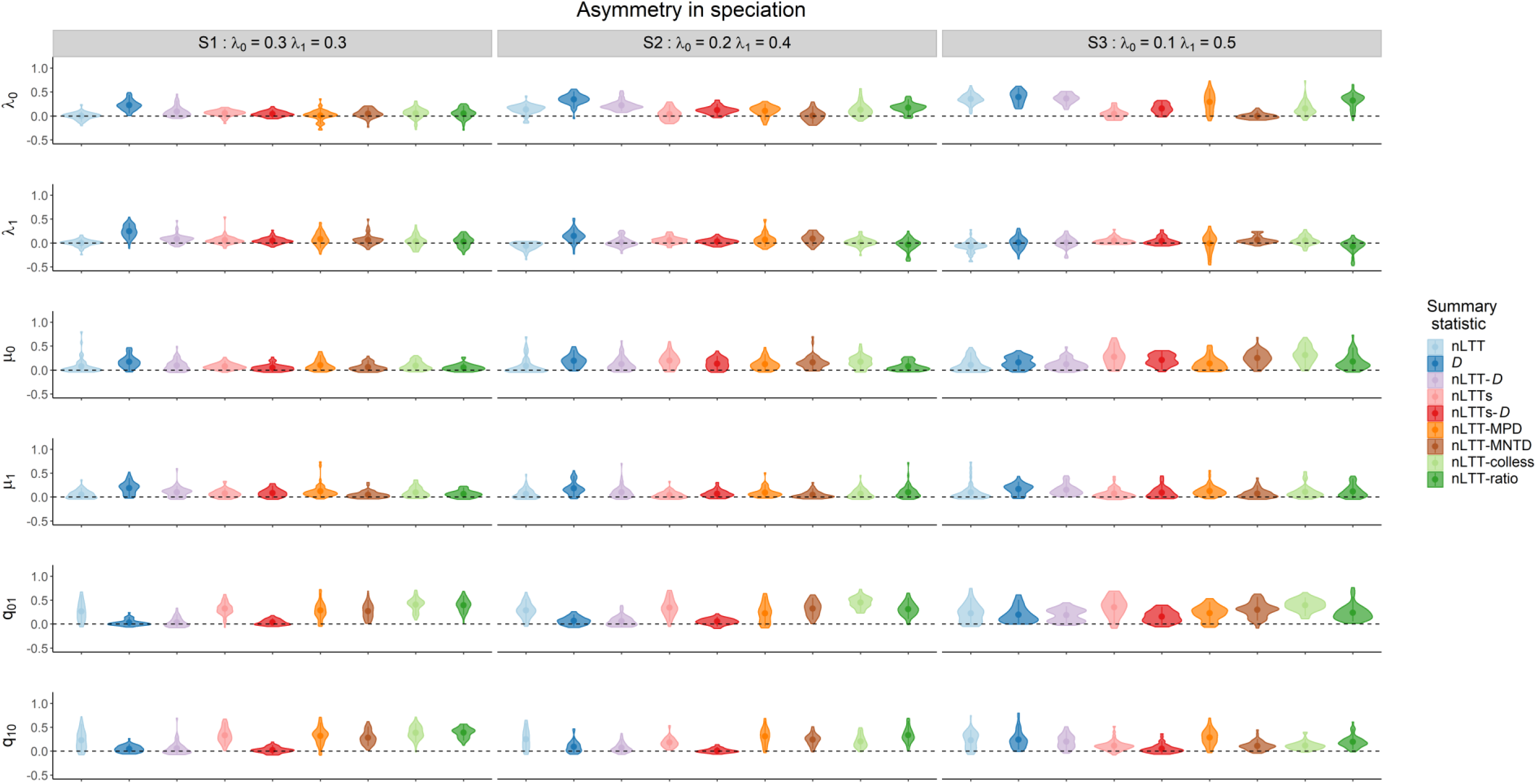
Parameter estimations of state-dependent speciation, extinction and transition rates using the ABC method with different **summary statistics** for scenarios with varying degrees of **asymmetry in speciation** (scenarios S1, S2, S3 in Table 1) in the generating rates. Plots show the residual inference error between estimated and (true) generated values. Dashed horizontal lines represent zero error to guide the eye. The colors indicate ABC results using different summary statistic combinations.

**Fig 4.**
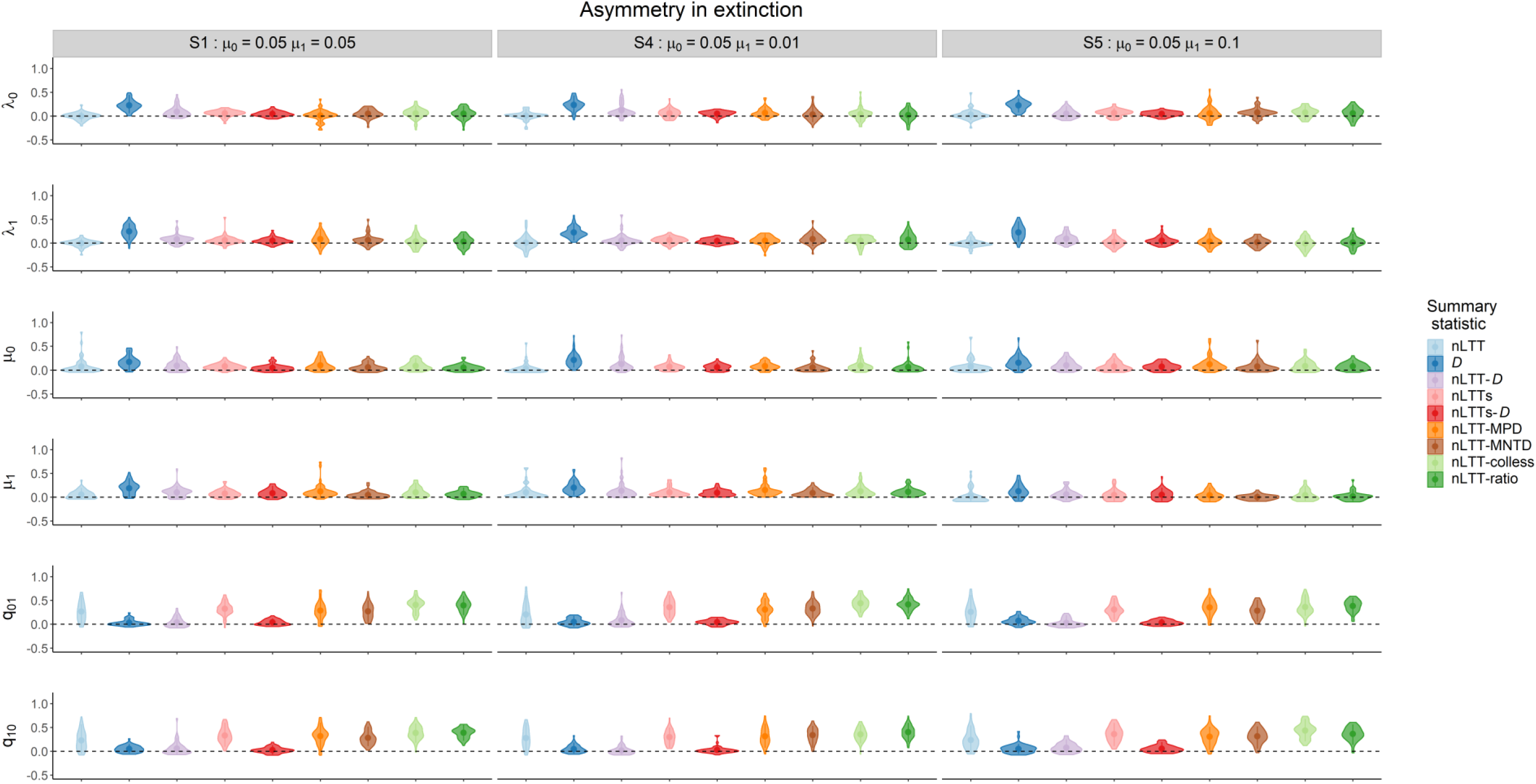
Parameter estimations of state-dependent speciation, extinction and transition rates using the ABC method with different **summary statistics** for scenarios with varying degrees of **asymmetry in extinction** (scenarios **S1, S4, S5** in Table 1) in the generating rates. Plots show the residual inference error between estimated and (true) generated values. Dashed horizontal lines represent zero error to guide the eye. The colors indicate ABC results using different summary statistic combinations.

**Fig 5.**
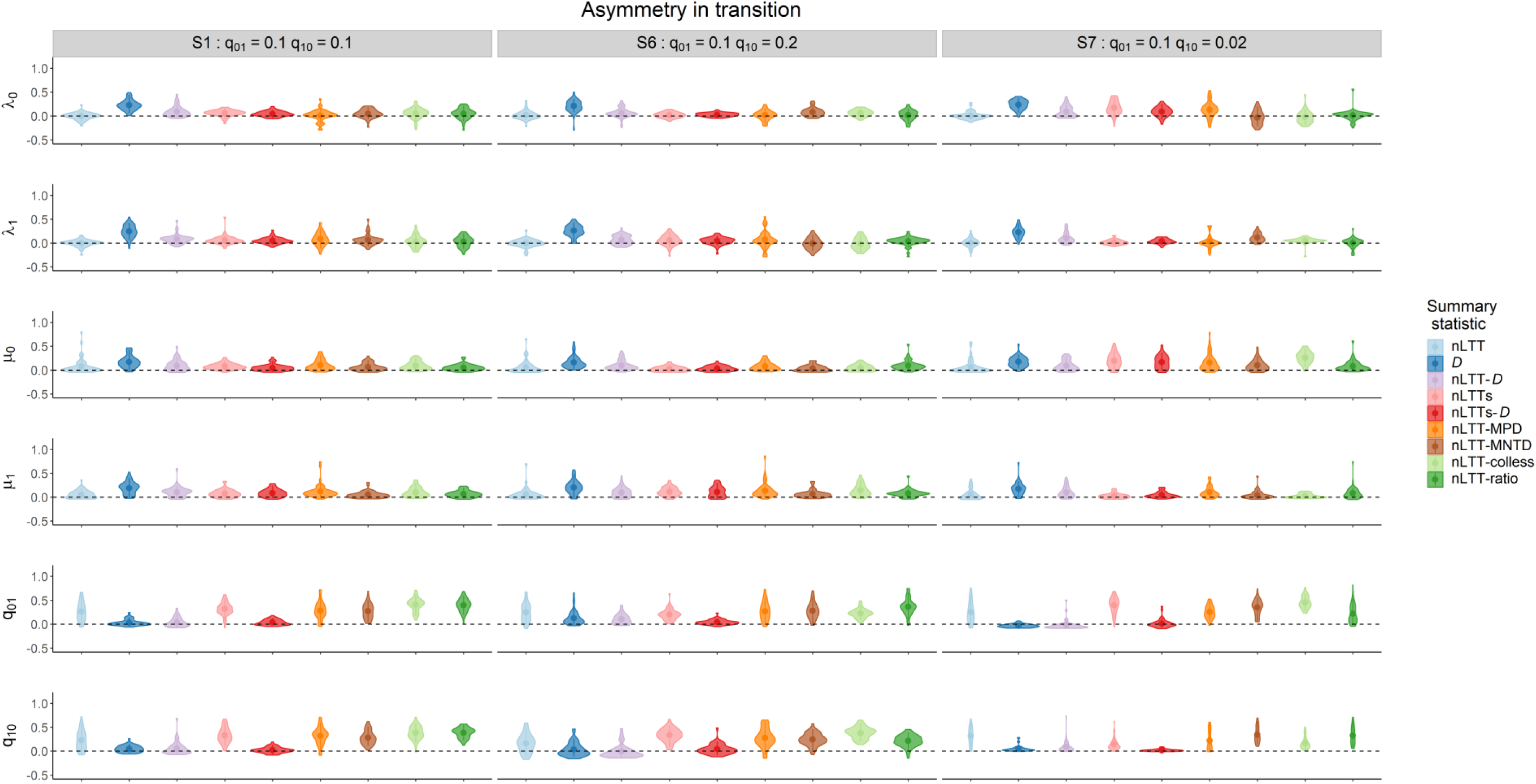
Parameter estimations of state-dependent speciation, extinction and transition rates using the ABC method with different **summary statistics** for scenarios with varying degrees of **asymmetry in transition** (scenarios **S1, S6, S7** in Table 1) in the generating rates. Plots show the residual inference error between estimated and (true) generated values. Dashed horizontal lines represent zero error to guide the eye. The colors indicate ABC results using different summary statistic combinations.

In contrast, the statistic *D* is efficient in estimating transition rates in different scenarios, but leads to large bias and variation in estimating speciation and extinction rates when used on its own (Figs 3 and 4). Combining the statistic *D* with the nLTT statistics (i.e., nLTT-*D* and nLTTs-*D*) visibly improves the estimation accuracy in transition rates in all the scenarios, as well as the diversification rates in extreme cases (tip ratio > 5) (Fig 3 and S1). Overall, the best summary statistic combination is nLTTs-*D*, which includes both trait dynamic information from phylogenetic signal (as measured by *D*) and temporal trait information coming from the nLTT statistics.

When evaluating the correlations between the summary statistics, we found that nLTT statistics have relatively strong positive or negative correlations with most of other summary statistics except *D*, especially a strong negative correlation with MNTD (Fig S4). This indicates that there is overlap of information among these summary statistics. Conversely, *D* is independent of most of the statistics, as it shows weak correlations (Fig S4).

### Inference with different methods (ABC, MCMC and MLE)

Of the nine summary statistic combinations we considered, the combination of the phylogenetic signal summary statistic *D* and the nLTT statistics (i.e., nLTTs-*D*) was found to give the most accurate state-dependent rate estimations (Figs 3, 4, 5 and S1). Therefore, here we focus on comparing the ABC estimations of nLTTs-*D* with the estimations of MCMC and MLE. Overall, the ABC method with efficient summary statistics performs well in estimating state-dependent rates, similar to the likelihood-based estimations in most scenarios. Zooming into the groups of scenarios with different levels of asymmetry in speciation, extinction and transition (Figs 6, 7 and 8), we found that asymmetry in speciation rates had a greater influence on inference accuracy than asymmetry in extinction or transition rates. In the scenarios with asymmetric speciation rates, the inference error increases with a higher level of asymmetry (Fig 6). But the bias only occurs when estimating the rates of the species-poor state (e.g., *λ_0_*, *μ_0_*, *q_01_* in scenario S3) (Fig 6). Similarly, when transition rates are highly asymmetric between states, the bias occurs in estimating extinction rate of the species-poor state (*μ_0_* in scenario S7) (Fig 8). The ABC method can always accurately estimate net diversification rates of each state in all the scenarios, with low bias and variance, even more so than the MCMC and MLE estimations in some scenarios (Fig S2).

**Fig 6.**
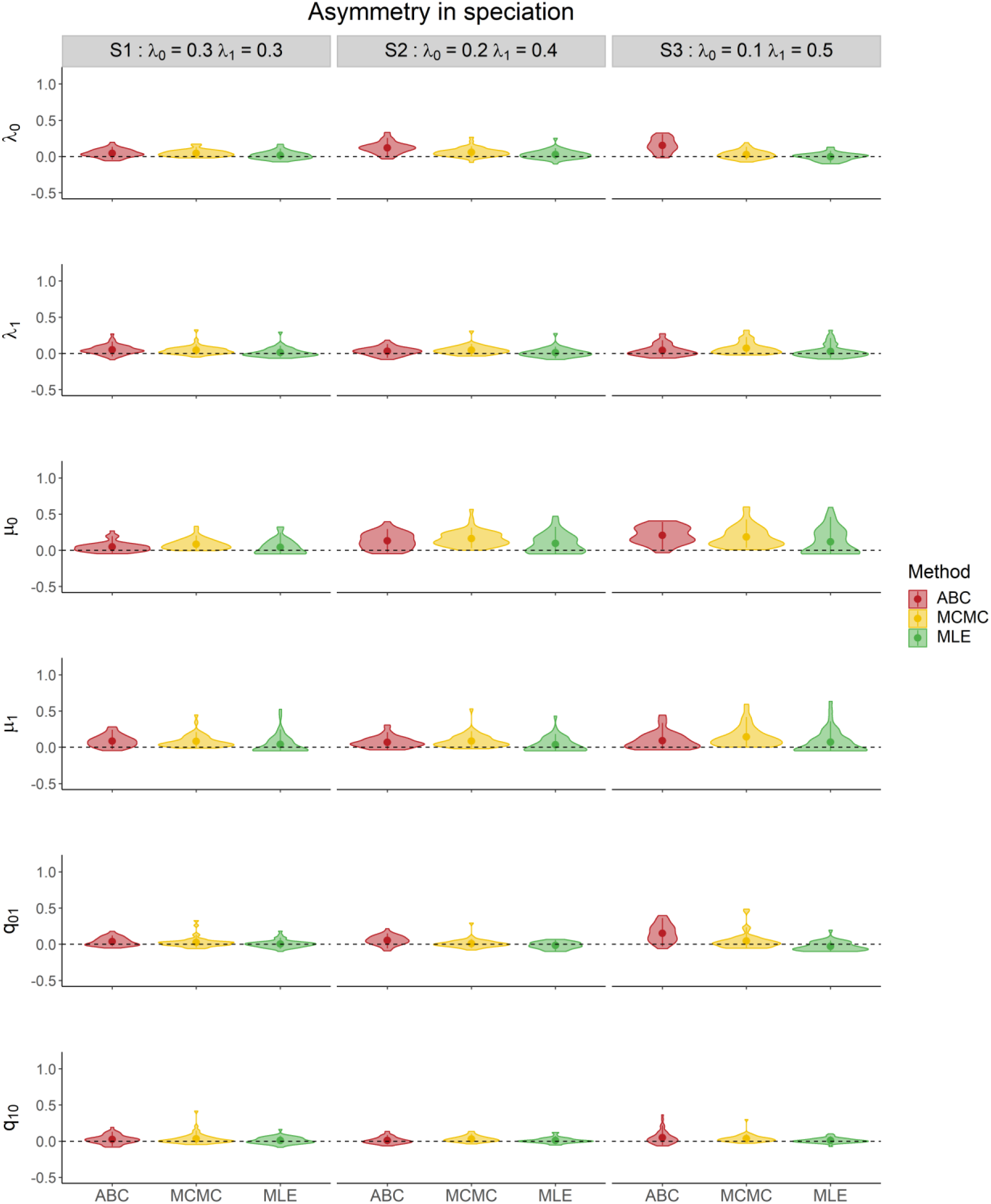
Parameter estimations of state-dependent speciation, extinction and transition rates using the **ABC, MCMC and MLE** methods for scenarios with varying degrees of **asymmetry in speciation** (scenarios S1, S2, S3 in Table 1) in the generation rates. Plots show the residual inference error between estimated and (true) generated values. Dashed horizontal lines represent zero error to guide the eye. Colors indicate different inference methods. The ABC results are estimated using the summary statistic combination nLTTs-*D* (nLTT_total_, nLTT_0_, nLTT_1_ and *D*).

**Fig 7.**
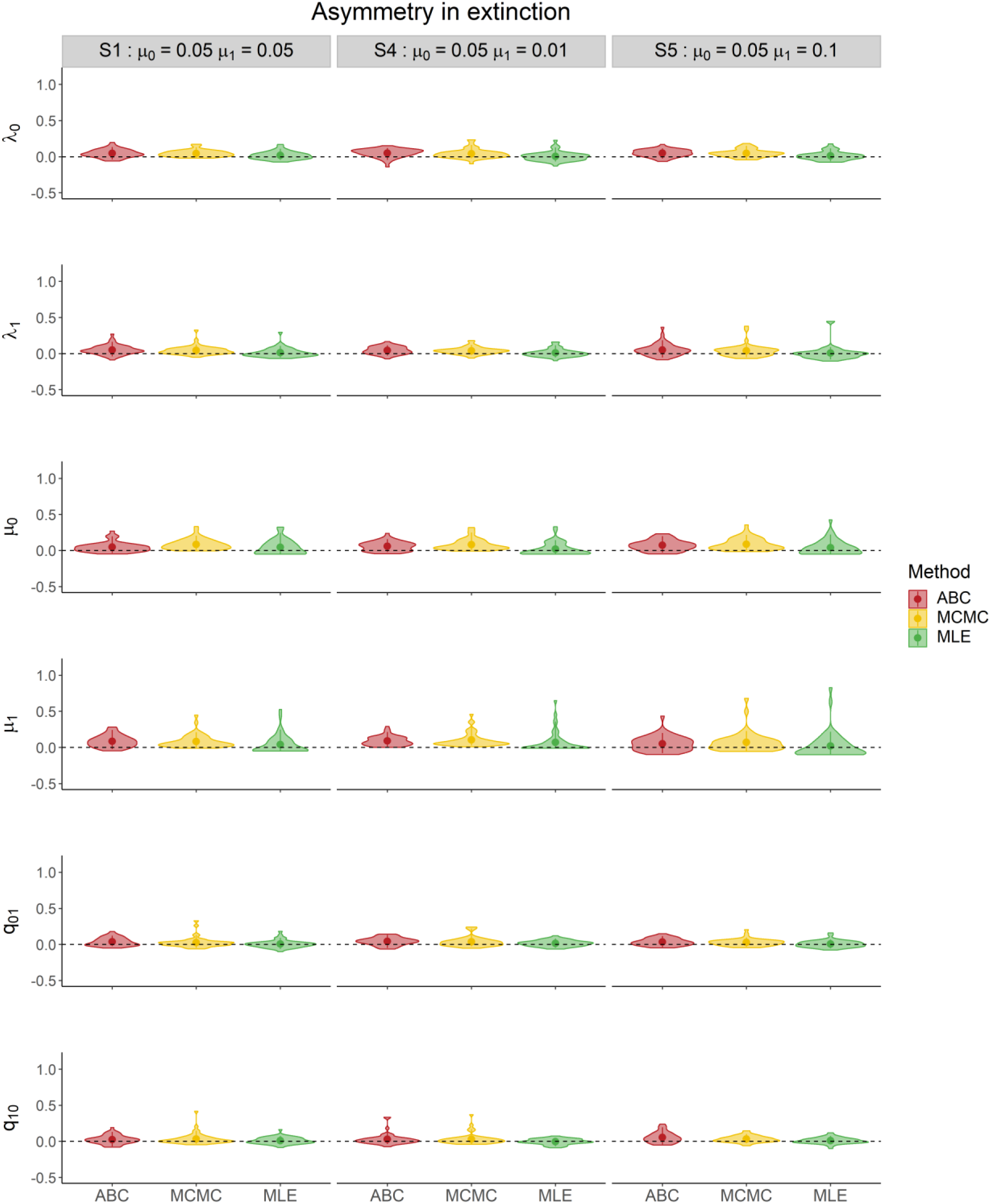
Parameter estimations of state-dependent speciation, extinction and transition rates using the **ABC, MCMC and MLE** methods for scenarios with varying degrees of **asymmetry in extinction** (scenarios S1, S4, S5 in Table 1) in the generating rates. Plots show the residual inference error between estimated and (true) generated values. Dashed horizontal lines represent zero error to guide the eye. Colors indicate different inference methods. The ABC results are estimated using the summary statistic combination nLTTs-*D* (nLTT_total_, nLTT_0_, nLTT_1_ and *D*).

**Fig 8.**
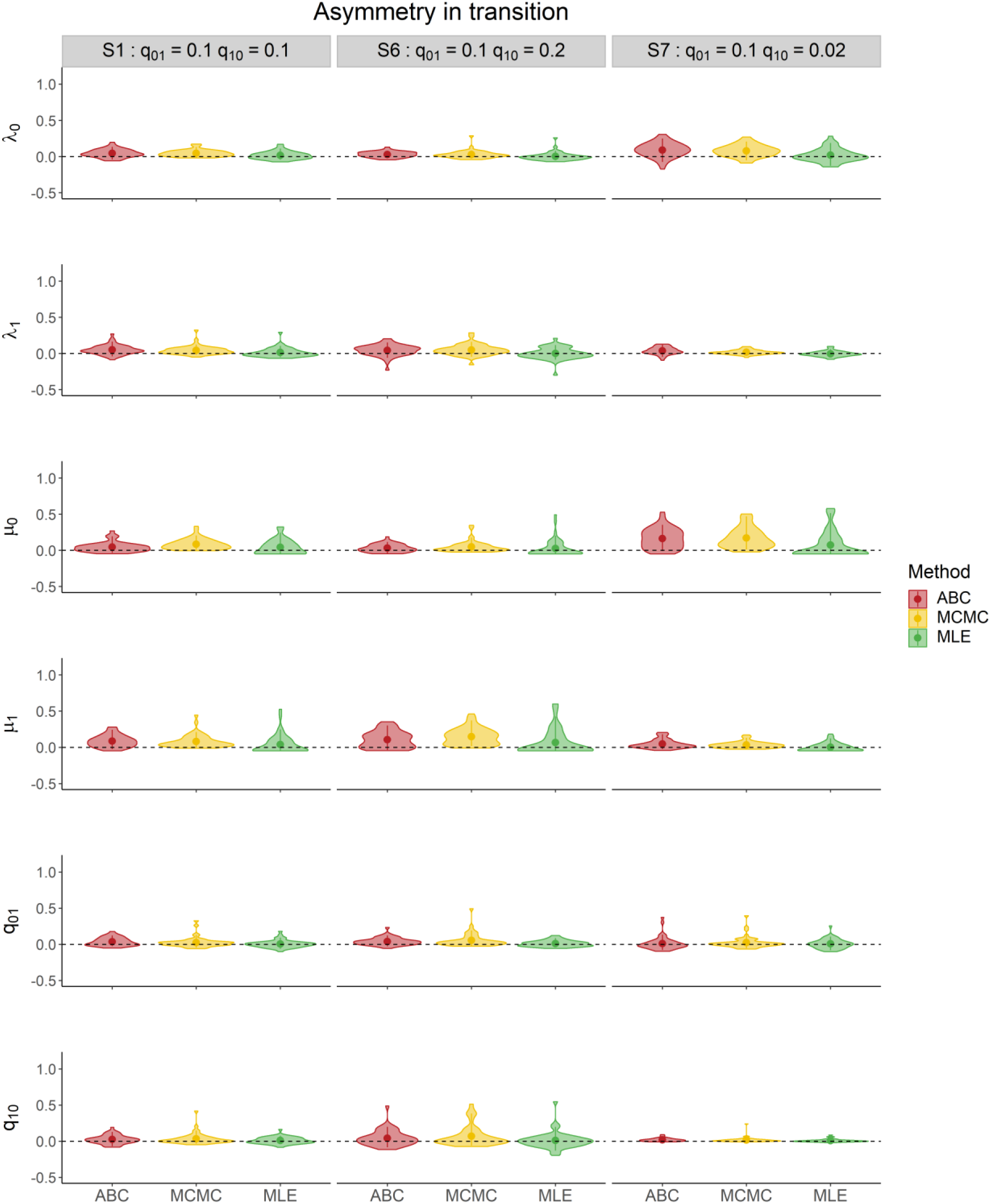
Parameter estimations of state-dependent speciation, extinction and transition rates using the **ABC, MCMC and MLE** methods for scenarios with varying degrees of **asymmetry in transition** (scenarios S1, S6, S7 in Table 1) in the generating rates. Plots show the residual inference error between estimated and (true) generated values. Dashed horizontal lines represent zero error to guide the eye. Colors indicate different inference methods. The ABC results are estimated using the summary statistic combination nLTTs-*D* (nLTT_total_, nLTT_0_, nLTT_1_ and *D*).

### Effect of statistic of observed data (tree size and tip ratio) on inference

We evaluated the relationships between tree size and tip ratio (Table 2) with inference error across all the observed datasets. As expected, small trees (< 100 tips) or trees with large diversity difference between states tend to cause larger inference error, especially in extinction rates, which in turn affect the estimation accuracy of the net diversification rates (Figs 9,10 and S5).

**Fig 9.**
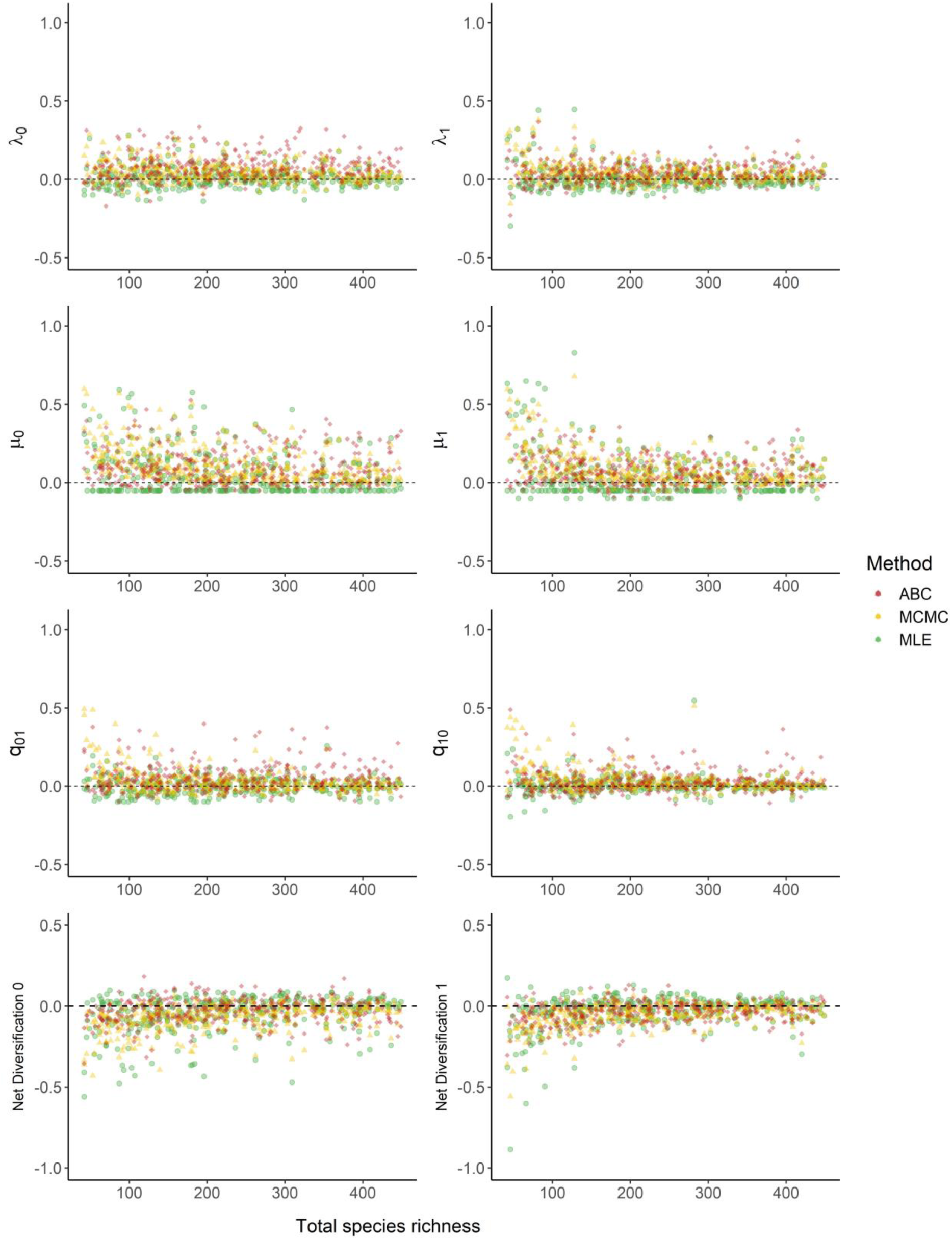
Relationship between **phylogenetic tree size** (total number of species) and the inference error in estimating state-dependent rates (estimated minus observed). The points show the median value of the posterior distribution in MCMC and ABC algorithms, and point estimates in MLE. Different color indicates different methods.

**Fig 10.**
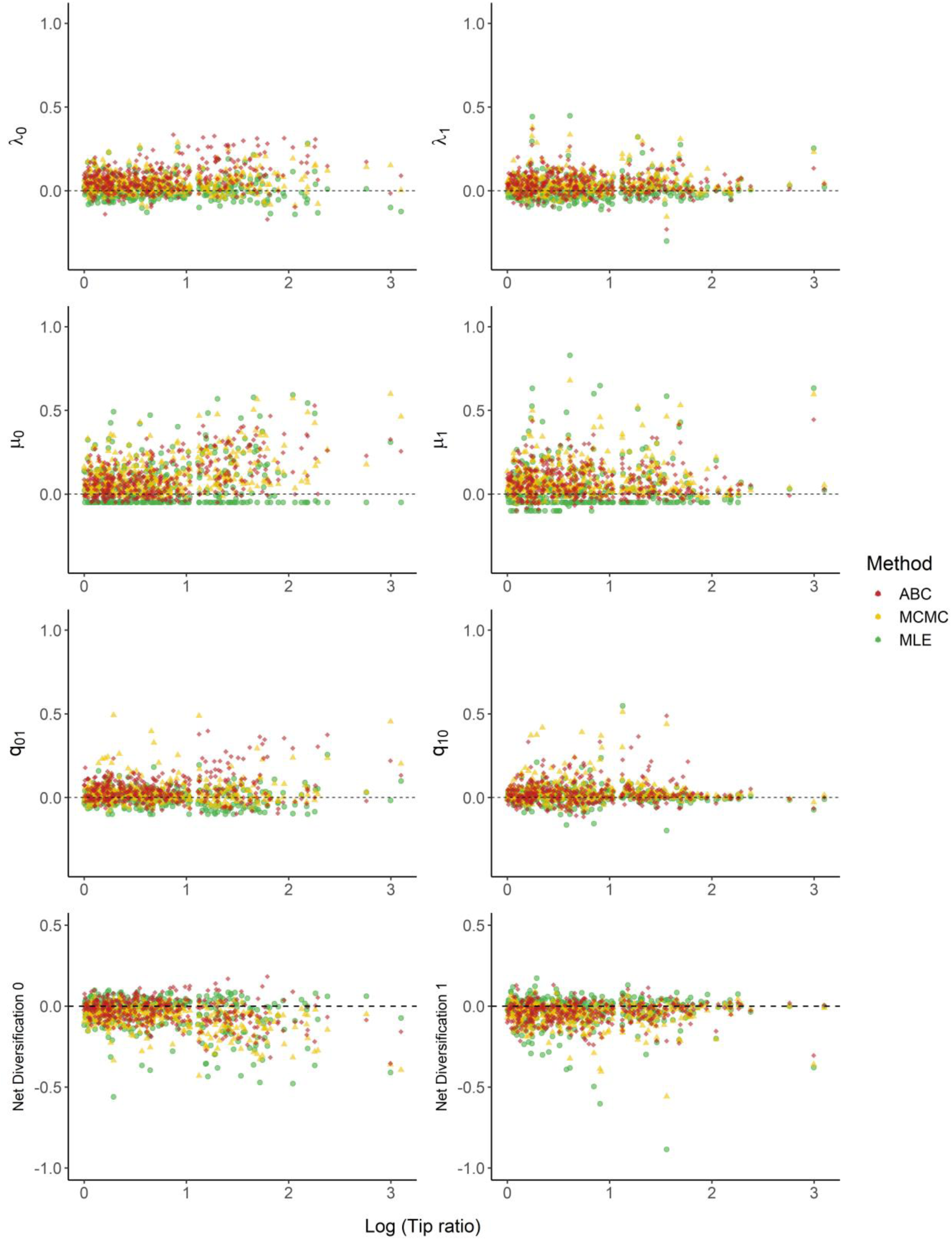
Relationship between the **tip ratio** and the inference error in estimating state-dependent rates. The points show the median value of the posterior distribution in MCMC and ABC algorithms, and point estimates in MLE. Different color indicates different methods.

## Discussion

The development of SSE models has recently accelerated, especially with advances in mathematical modelling techniques and the availability of empirical data (Holland et al., 2020). The main aim of this study was not to evaluate the power of existing likelihood methods of SSE models, which has already been done in previous studies (Davis et al., 2013; Holland et al., 2020). Instead, we used the likelihood-based estimations as a baseline to evaluate the performance of a new likelihood-free ABC approach, and to search for efficient summary statistics that cover the most comprehensive phylogenetic and trait information. Across all the scenarios we tested, the combination of the nLTT statistics and phylogenetic signal *D* is sufficient to produce accurate estimations in ABC, which is on par with the likelihood estimations.

The nLTT statistic, which provides information of evolutionary dynamics over time, has been shown to be efficient and informative in phylogenetic analysis and diversification studies (Janzen et al., 2015; Saulnier et al., 2017; Richter & Wit, 2021), as well in island biogeography (Xie et al., 2023). However, the power of the statistic is limited in trait-related analysis when estimating transition rates, due to the challenge and uncertainty of mapping trait evolution on phylogenies. nLTT for the whole tree can only capture the average diversification rates independent of traits, while adding nLTT for each state (nLTT_0_ and nLTT_1_) can improve the ability in capturing trait dependency in speciation and extinction rates (Figs 3), as well as net diversification rates (Fig S1), because of the information of branching times in the reduced trees. It is similar to the original sister-clade comparison method, which has been widely used in detecting the effect of traits on diversification rates before the establish of the SSE models (Mitter et al., 1988; Maddison et al., 2007). Therefore, likewise, nLTT has the same limitation on detecting state shifts over time, leading to a poor estimation of transition rates between states (Figs 3, 4 and 5). However, the phylogenetic signal statistic *D* contains efficient information on trait evolution, by comparing the observed trait distribution with the distributions simulated through continuous Brownian Motion patterns, compensating for the lack of the information in nLTT statistics.

In this paper, we only tested the performance of nine combinations among six summary statistics. However, there are a number of alternative statistics available in phylogenetics (e.g., phylogenetic diversity (*PD*), *Laplacian spectrum* for tree shape). Recently, a few of studies have proposed a number of summary statistics for detecting trait dynamics. Lajaaiti et al. (2023) applied multiple neural network architectures to estimate state-dependent diversification rates, and compared the performance of the deep learning methods with the maximum likelihood estimation, which provided a promising alternative for phylogenetic inference. However, the study focuses more on summary statistics for describing the shape of phylogenetic trees (84 summary statistics), but less so on trait dynamics (one summary statistic (tip ratio)). Thereafter, Schwery et al., (2023) used a Bayesian approach to test the adequacy of trait-dependent diversification models with a number of summary statistics, selected to capture trait distributions (e.g., FiSSE statistics etc.) and the features of the phylogenetic tree (e.g., gamma statistics, Colless index, branch length, etc.). A few of summary statistics that have not been included in our study are worth testing, but that does not mean adding more summary statistic is necessarily better or recommended. A primary motivation for using summary statistics is to reduce the dimensionality of the observed datasets, whilst retaining the most information for parameter estimation. Selecting an excess number of summary statistics, especially those that are highly correlated with one another, may lead to overfitting and redundancy, which reduce the efficiency and computability of the ABC algorithm (Jung & Marjoram, 2011; Blum et al., 2013). Therefore, apart from looking for alternative summary statistics, it is also important to choose and construct efficient combinations by weighting or transforming the statistics.

A number of methods have been developed to address the challenges of identifying appropriate summary statistics (Wegmann et al., 2009; Nunes & Balding, 2010; Fearnhead & Prangle, 2012). Joyce and Marjoram (2008) introduced a sequential scheme to choose a sufficient subset of summary statistics by adding a randomly chosen statistic in each iteration, and evaluating whether the inclusion of the additional statistic improves the inference ability. However, the drawback is that the selected subset depends on the order of the additional summary statistics, that is, when more informative summary statistics are added late and less informative statistics have been included, the final statistic combination may be found to be redundant. Jung and Marjoram (2011) improved the method by assigning weights to each summary statistic. The method keeps all the statistics rather than filtering from the statistic pool, and allows higher weights to the statistics that are more informative. In addition, Wegmann et al., (2009) introduced a statistical approach using a partial least squares (PLS) regression, which is powerful to reduce the dimensionality of the variables. The method extracts the orthogonal components as a subset of informative summary statistics, which are highly corelated with the parameters but decorrelated with each other. Later, a semi-automatic procedure was proposed in ABC algorithms to construct summary statistics by reducing dimension with a regression-based approach, which improves both the performance of parameter estimation and model selection (Fearnhead & Prangle, 2012; Prangle et al., 2014; Harrison Id & Baker, 2020). These methods are more effective in filtering efficient summary statistics than manual selection, and may provide great insights for further improvements of ABC efficiency.

We found that small observed trees lead to high inference error in BiSSE estimation using both MLE, MCMC and ABC methods, especially in estimating extinction rates (Figs 9 and S5), which reemphasizes the results of previous studies that explored the accuracy of power of SSE models (Davis et al., 2013; Gamisch, 2016). Similarly, in ABC estimations, large variance in speciation rates (*λ*_1_ / *λ*_0_ = 5) or transition rates (q_01_ / q_10_ = 5) leads to a limited number of existing species in one state. Therefore, the summary statistics using trimmed trees (with either state) for trait analysis cause inference error in the specie-poor state, due to the lack of information of phylogenetic and trait dynamics in this state. On the other hand, the inference error in the extreme scenario may be due to the constraint of the total number of species (< 500 species in simulated trees) in our study. To test the effects of the constraint, we run each scenario for 1000 replicates without constraints, and calculate the proportion of the replicates that contain fewer than 500 species, the results of which are shown in Table 3. The extreme case (scenario S3) that we are most interested in (strong asymmetry in speciation) is the most affected by the constraint, which means our sample of observed data is biased. The constraint results in more sampled observed phylogenies with ancestral species under the lower-rate state. These phylogenetic trees have a slow diversification at early stage until some branches transition to the higher-rate state, which may lead to a late burst before present time. We note that currently the ABC method may fail to determine whether the poor richness of a certain state is due to a low speciation rate or a high transition rate to the other state in this case (Fig S5).

**Table 3.** Proportion of the replicates with fewer than 500 species in 1000 randomly sampled trees.

| Scenario | Speciation<br>rate 0 ($\lambda_0$) | Speciation<br>rate 1 ($\lambda_1$) | Extinction<br>rate 0 ($\mu_0$) | Extinction<br>rate 1 ($\mu_1$) | Transition<br>0 to 1<br>($q_{01}$) | Transition<br>1 to 0<br>($q_{10}$) | Proportion<br>replicates<br><500<br>species |
| --- | --- | --- | --- | --- | --- | --- | --- |
| S1 | 0.3 | 0.3 | 0.05 | 0.05 | 0.1 | 0.1 | 0.94 |
| S2 | 0.2 | 0.4 | 0.05 | 0.05 | 0.1 | 0.1 | 0.71 |
| S3 | 0.1 | 0.5 | 0.05 | 0.05 | 0.1 | 0.1 | 0.46 |
| S4 | 0.3 | 0.3 | 0.05 | 0.01 | 0.1 | 0.1 | 0.86 |
| S5 | 0.3 | 0.3 | 0.05 | 0.1 | 0.1 | 0.1 | 0.99 |
| S6 | 0.3 | 0.3 | 0.05 | 0.05 | 0.1 | 0.2 | 0.95 |
| S7 | 0.3 | 0.3 | 0.05 | 0.05 | 0.1 | 0.02 | 0.94 |

Apart from the constraint of the size of datasets, another typical issue with the power of the BiSSE model is high type I error (Rabosky & Goldberg, 2015). This problem has been solved in a more complex model HiSSE and derived models. Currently we have only developed the ABC estimation in the BiSSE model, but it can be easily extended to more complex models, such as SecSSE, especially since our simulations have been generated with the *secsse* package. Furthermore, machine learning, as a rapidly developing methodology, has been incorporated into the ABC algorithms (Mondal et al., 2019; Sanchez et al., 2020), and may have good potential for future studies. In any case, both ABC and machine learning methods offer important opportunities for further expansions of SSE models incorporating more factors that affect evolutionary patterns (e.g., geographical and ecological factors), where the likelihood equations are too complex to solve.

## Reference

Adams, D. C. (2013). Comparing evolutionary rates for different phenotypic traits on a phylogeny using likelihood. Systematic Biology, 62(2), 181–192. 10.1093/sysbio/sys083

Armbruster, W. S. (2014). Floral specialization and angiosperm diversity: Phenotypic divergence, fitness trade-offs and realized pollination accuracy. AoB PLANTS, 6, 1–24. 10.1093/aobpla/plu003

Bartoszek, K., & Liò, P. (2019). Modelling trait-dependent speciation with approximate Bayesian computation. *Acta Physica Polonica B*, Proceedings Supplement, 12(1), 25–47. 10.5506/APhysPolBSupp.12.25

Beaulieu, J. M., & O’Meara, B. C. (2016). Detecting hidden diversification shifts in models of trait-dependent speciation and extinction. Systematic Biology, 65(4), 583–601. 10.1093/sysbio/syw022

Beaumont, M. A. (2010). Approximate Bayesian Computation in Evolution and Ecology. Annu. Rev. Ecol. Evol. Syst, 41, 379–406. 10.1146/annurev-ecolsys-102209-144621

Beaumont, M. A. (2019). Approximate Bayesian computation. Annual Review of Statistics and Its Application, 6, 379–403. 10.1146/annurev-statistics-030718-105212

Beaumont, M. A., Cornuet, J. M., Marin, J. M., & Robert, C. P. (2009). Adaptive approximate Bayesian computation. Biometrika, 96(4), 983–990. 10.1093/biomet/asp052

Blomberg, S. P., Garland, T., & Ives, A. R. (2003). TESTING FOR PHYLOGENETIC SIGNAL IN COMPARATIVE DATA: BEHAVIORAL TRAITS ARE MORE LABILE. Evolution, 57(4), 717–745. 10.1111/J.0014-3820.2003.TB00285.X

Blum, M. G. B., Nunes, M. A., Prangle, D., & Sisson, S. A. (2013). A comparative review of dimension reduction methods in approximate bayesian computation. Statistical Science, 28(2), 189–208. 10.1214/12-STS406

Borges, R., Machado, J. P., Gomes, C., Rocha, A. P., & Antunes, A. (2019a). Measuring phylogenetic signal between categorical traits and phylogenies. Bioinformatics, 35(11), 1862–1869. 10.1093/BIOINFORMATICS/BTY800

Borges, R., Machado, J. P., Gomes, C., Rocha, A. P., & Antunes, A. (2019b). Measuring phylogenetic signal between categorical traits and phylogenies. Bioinformatics, 35(11), 1862–1869. 10.1093/BIOINFORMATICS/BTY800

Colless, D. H. (1995). Relative Symmetry of Cladograms and Phenograms: An Experimental Study. Systematic Biology, 44(1), 102–108. 10.1093/SYSBIO/44.1.102

Csilléry, K., Blum, M. G. B., Gaggiotti, O. E., & François, O. (2010). Approximate Bayesian Computation (ABC) in practice. Trends in Ecology and Evolution, 25(7), 410–418. 10.1016/j.tree.2010.04.001

Davis, M. P., Midford, P. E., & Maddison, W. (2013). Exploring power and parameter estimation of the BiSSE method for analyzing species diversification. BMC Evolutionary Biology, 13(1). 10.1186/1471-2148-13-38

Fearnhead, P., & Prangle, D. (2012a). Constructing summary statistics for approximate Bayesian computation: semi-automatic approximate Bayesian computation. J. R. Statist. Soc. B, 74, 419–474.

Fearnhead, P., & Prangle, D. (2012b). Constructing summary statistics for approximate Bayesian computation: Semi-automatic approximate Bayesian computation. Journal of the Royal Statistical Society. Series B: Statistical Methodology, 74(3), 419–474. 10.1111/j.1467-9868.2011.01010.x

Fernández-Mazuecos, M., Blanco-Pastor, J. L., Juan, A., Carnicero, P., Forrest, A., Alarcón, M., Vargas, P., & Glover, B. J. (2019). Macroevolutionary dynamics of nectar spurs, a key evolutionary innovation. The New Phytologist, 222(2), 1123–1138. 10.1111/NPH.15654

Fitzjohn, R. G. (2010). Quantitative traits and diversification. Systematic Biology, 59(6), 619–633. 10.1093/sysbio/syq053

Fitzjohn, R. G. (2012). Diversitree: Comparative phylogenetic analyses of diversification in R. Methods in Ecology and Evolution, 3(6), 1084–1092. 10.1111/j.2041-210X.2012.00234.x

FitzJohn, R. G. (2012). Diversitree: comparative phylogenetic analyses of diversification in R. Methods in Ecology and Evolution, 3(6), 1084–1092. 10.1111/J.2041-210X.2012.00234.X

Fritz, S. A., & Purvis, A. (2010). Selectivity in mammalian extinction risk and threat types: A new measure of phylogenetic signal strength in binary traits. Conservation Biology, 24(4), 1042–1051. 10.1111/j.1523-1739.2010.01455.x

Gamisch, A. (2016). Notes on the Statistical Power of the Binary State Speciation and Extinction (BiSSE) Model. Evolutionary Bioinformatics Online, 12, 165–174. 10.4137/EBO.S39732

Goldberg, E. E., Lancaster, L. T., & Ree, R. H. (2011). Phylogenetic inference of reciprocal effects between geographic range evolution and diversification. Systematic Biology, 60(4), 451–465. 10.1093/sysbio/syr046

Harrison Id, J. U., & Baker, R. E. (2020). An automatic adaptive method to combine summary statistics in approximate Bayesian computation. 10.1371/journal.pone.0236954

Herrera-Alsina, L., Van Els, P., & Etienne, R. S. (2019). Detecting the Dependence of Diversification on Multiple Traits from Phylogenetic Trees and Trait Data. Systematic Biology, 68(2), 317–328. 10.1093/sysbio/syy057

Holland, B. R., Ketelaar-Jones, S., O’Mara, A. R., Woodhams, M. D., & Jordan, G. J. (2020). Accuracy of ancestral state reconstruction for non-neutral traits. Scientific Reports, 10(1), 1–10. 10.1038/s41598-020-64647-4

Janzen, T., Alzate, A., Muschick, M., Maan, M. E., van der Plas, F., & Etienne, R. S. (2017). Community assembly in Lake Tanganyika cichlid fish: quantifying the contributions of both niche-based and neutral processes. Ecology and Evolution, 7(4), 1057–1067. 10.1002/ece3.2689

Janzen, T., Höhna, S., & Etienne, R. S. (2015). Approximate Bayesian Computation of diversification rates from molecular phylogenies: introducing a new efficient summary statistic, the nLTT. Methods in Ecology and Evolution, 6(5), 566–575. 10.1111/2041-210X.12350

Joyce, P., & Marjoram, P. (2008). Approximately sufficient statistics and bayesian computation. Statistical Applications in Genetics and Molecular Biology, 7(1). 10.2202/1544-6115.1389

Jung, H., & Marjoram, P. (2011). Choice of summary statistic weights in approximate bayesian computation. Statistical Applications in Genetics and Molecular Biology, 10(1). 10.2202/1544-6115.1586

Laudanno, G., Haegeman, B., Rabosky, D. L., & Etienne, R. S. (2021). Detecting Lineage-Specific Shifts in Diversification: A Proper Likelihood Approach. Systematic Biology, 70(2), 389–407. 10.1093/sysbio/syaa048

Li, P., & Wiens, J. J. (2022). What drives diversification? Range expansion tops climate, life history, habitat and size in lizards and snakes. Journal of Biogeography, 49(2), 237–247. 10.1111/JBI.14304

Louca, S., & Pennell, M. W. (2020). A General and Efficient Algorithm for the Likelihood of Diversification and Discrete-Trait Evolutionary Models. Systematic Biology, 69(3), 545–556. 10.1093/sysbio/syz055

Maddison, W. P., Midford, P. E., & Otto, S. P. (2007). Estimating a binary character’s effect on speciation and extinction. Systematic Biology, 56(5), 701–710. 10.1080/10635150701607033

Marjoram, P., Molitor, J., Plagnol, V., & Tavaré, S. (2003). Markov chain Monte Carlo without likelihoods. Proceedings of the National Academy of Sciences of the United States of America, 100(26), 15324–15328. 10.1073/pnas.0306899100

Miles, D. B., & Dunham, A. E. (1993). Historical perspectives in ecology and evolutionary biology: The use of phylogenetic comparative analyses. Annual Review of Ecology and Systematics, 24(February 2016), 587–619. 10.1146/annurev.es.24.110193.003103

Mitter, C., Farrell, B., & Wiegmann, B. (1988). The phylogenetic study of adaptive zones: has phytophagy promoted insect diversification? American Naturalist, 132(1), 107–128. 10.1086/284840

Mondal, M., Bertranpetit, J., & Lao, O. (2019). Approximate Bayesian computation with deep learning supports a third archaic introgression in Asia and Oceania. 10.1038/s41467-018-08089-7

Nunes, M. A., & Balding, D. J. (2010). On optimal selection of summary statistics for approximate bayesian computation. Statistical Applications in Genetics and Molecular Biology, 9(1). 10.2202/1544-6115.1576

Onstein, R. E., Baker, W. J., Couvreur, T. L. P., Faurby, S., Svenning, J. C., & Kissling, W. D. (2017). Frugivory-related traits promote speciation of tropical palms. Nature Ecology & Evolution 2017 1:12, 1(12), 1903–1911. 10.1038/s41559-017-0348-7

Pagel, M. (1999). Inferring the historical patterns of biological evolution. Nature, 401(6756), 877–884. 10.1038/44766

Prangle, D., Fearnhead, P., Cox, M. P., Biggs, P. J., & French, N. P. (2014). Semi-automatic selection of summary statistics for ABC model choice. Statistical Applications in Genetics and Molecular Biology, 13(1), 67–82. 10.1515/sagmb-2013-0012

Pyron, R. A., & Burbrink, F. T. (2012). Trait-dependent diversification and the impact of palaeontological data on evolutionary hypothesis testing in New World ratsnakes (tribe Lampropeltini). Journal of Evolutionary Biology, 25(3), 497–508. 10.1111/j.1420-9101.2011.02440.x

Rabosky, D. L., & Goldberg, E. E. (2015). Model inadequacy and mistaken inferences of trait-dependent speciation. Systematic Biology, 64(2), 340–355. 10.1093/sysbio/syu131

Richter, F., & Wit, E. C. (2021). D ETECTING PHYLODIVERSITY - DEPENDENT DIVERSIFICATION WITH A GENERAL PHYLOGENETIC INFERENCE FRAMEWORK. 1–19.

Rolland, J., Condamine, F. L., Jiguet, F., & Morlon, H. (2014). Faster Speciation and Reduced Extinction in the Tropics Contribute to the Mammalian Latitudinal Diversity Gradient. PLoS Biology, 12(1). 10.1371/JOURNAL.PBIO.1001775

Sanchez, T., Cury, J., Charpiat, G., & Jay, F. (2020). Deep learning for population size history inference: Design, comparison and combination with approximate Bayesian computation. Molecular Ecology Resources, January, 1–16. 10.1111/1755-0998.13224

Saulnier, E., Gascuel, O., & Alizon, S. (2017). Inferring epidemiological parameters from phylogenies using regression-ABC: A comparative study. In PLoS Computational Biology (Vol. 13, Issue 3). 10.1371/journal.pcbi.1005416

Schwery, O., Freyman, W., & Goldberg, E. E. (2023). adequaSSE: Model Adequacy Testing for Trait-Dependent Diversification Models. 10.1101/2023.03.06.531416

Sirén, J., & Kaski, S. (2020). Local dimension reduction of summary statistics for likelihood-free inference. Statistics and Computing, 30(3), 559–570. 10.1007/S11222-019-09905-W/FIGURES/4

Tavare, S., Balding, D. J., Griffiths, J. R. C., & Donneuyst, P. (1997). From DNA. Genetics, 518(2), 505. http://www.genetics.org/cgi/content/abstract/145/2/505

Toni, T., Welch, D., Strelkowa, N., Ipsen, A., & Stumpf, M. P. H. (2009). Approximate Bayesian computation scheme for parameter inference and model selection in dynamical systems. Journal of the Royal Society Interface, 6(31), 187. 10.1098/RSIF.2008.0172

Vasconcelos, T., O’Meara, B. C., & Beaulieu, J. M. (2022). A flexible method for estimating tip diversification rates across a range of speciation and extinction scenarios. Evolution, 76(7), 1420–1433. 10.1111/evo.14517

Webb, C. O. (2000). Exploring the phylogenetic structure of ecological communities: An example for rain forest trees. American Naturalist, 156(2), 145–155. 10.1086/303378

Wegmann, D., Leuenberger, C., & Excoffier, L. (2009). Efficient approximate Bayesian computation coupled with Markov chain Monte Carlo without likelihood. Genetics, 182(4), 1207–1218. 10.1534/genetics.109.102509

Wiens, J. J. (2017). What explains patterns of biodiversity across the Tree of Life?: New research is revealing the causes of the dramatic variation in species numbers across branches of the Tree of Life. BioEssays, 39(3), 1–10. 10.1002/bies.201600128

Xu, L., Van Doorn, S., Hildenbrandt, H., & Etienne, R. S. (2021). Inferring the Effect of Species Interactions on Trait Evolution. Syst. Biol, 70(3), 463–479. 10.1093/sysbio/syaa072

