## Supplemental Figures for "Inferring state-dependent diversification rates using approximate Bayesian computation (ABC)"

**Supplementary Materials**

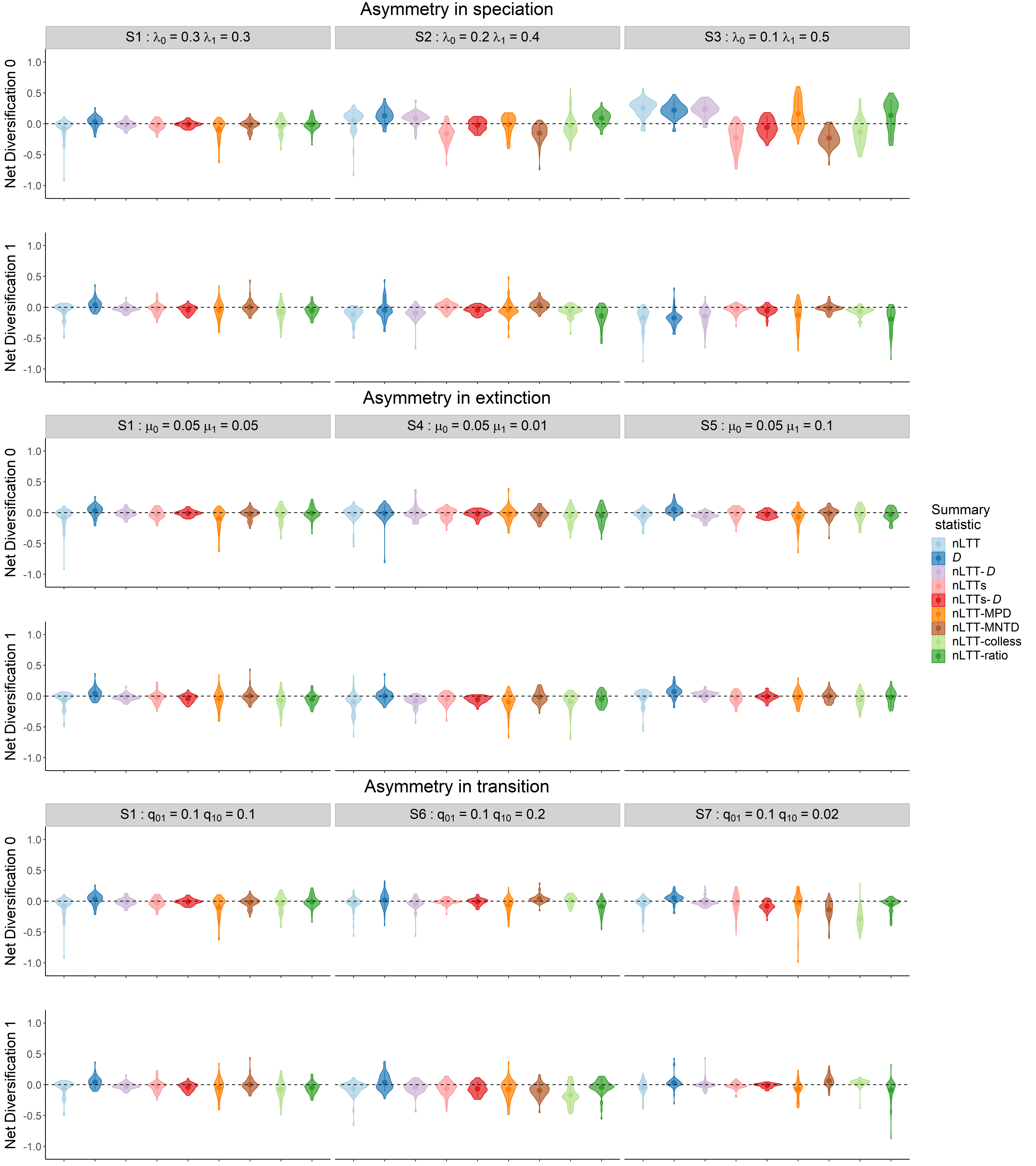

Figure S1. Parameter estimations of state-dependent **net diversification** (speciation - extinction) rates using the ABC method with different **summary statistics** when the generating rates have varying degrees of **asymmetry in** **speciation** (scenarios S1, S2, S3 in Table 1), **extinction** (scenarios S1, S4, S5 in Table 1), **transition** (scenarios S1, S6, S7 in Table 1). Plots show the residual inference error between estimated and (true) generated values. Dashed horizontal lines represent zero error to guide the eye. Each distribution combines 50 replicates for each scenario. The colors indicate ABC results with different summary statistic combinations.

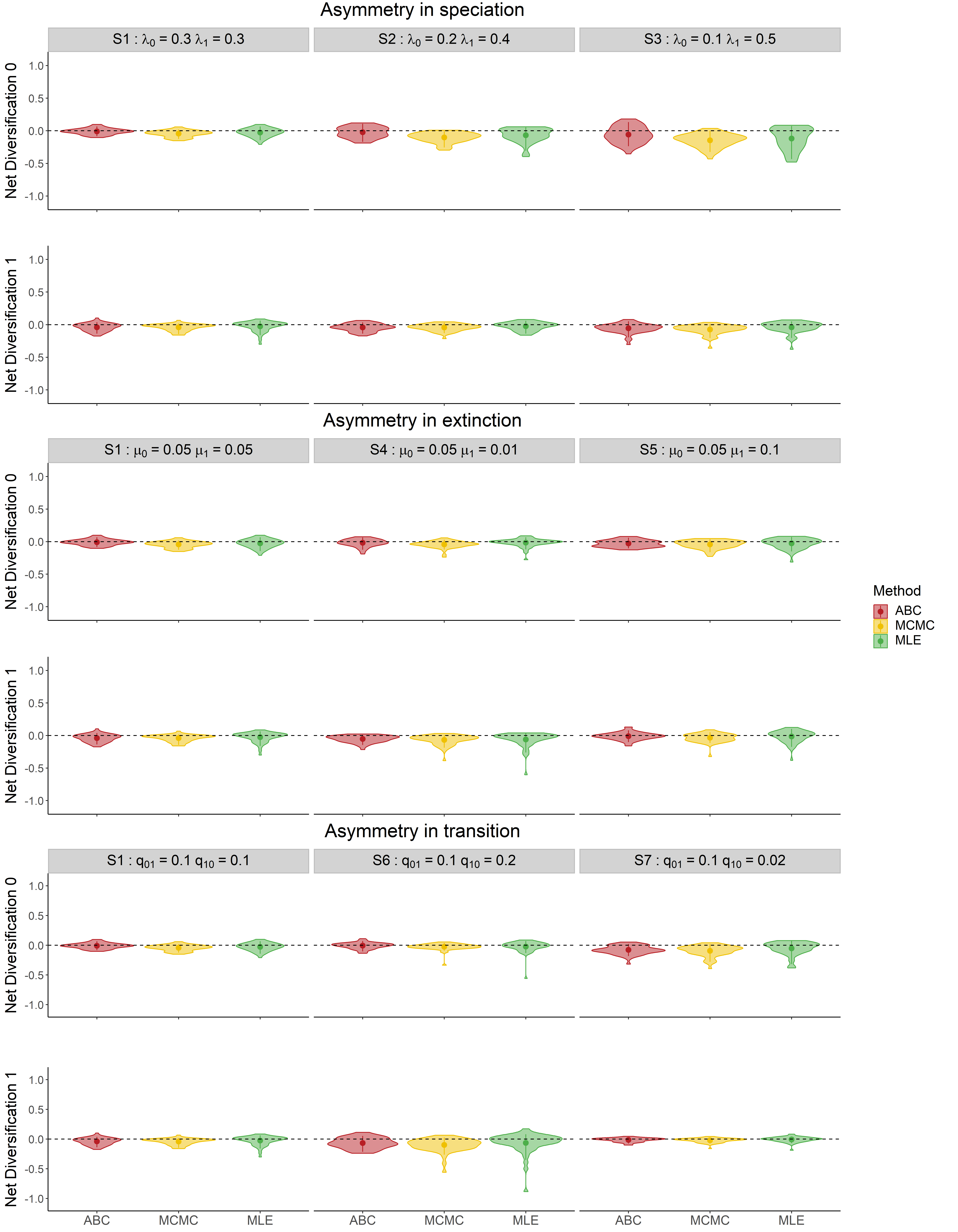

Figure S2. Parameter estimations of state-dependent **net diversification** (speciation - extinction) rates using the ABC, MCMC and MLE methods, when the generating rates have varying degrees of **asymmetry in** **speciation** (scenarios S1, S2, S3 in Table 1), **extinction** (scenarios S1, S4, S5 in Table 1), **transition** (scenarios S1, S6, S7 in Table 1). The colors indicate different inference methods. The plot shows the residual inference error after subtracting the generating values (estimation - generation). The ABC results are estimated using the summary statistic combination nLTTs-*D* (nLTT_total_, nLTT_0_, nLTT_1_ and *D*).

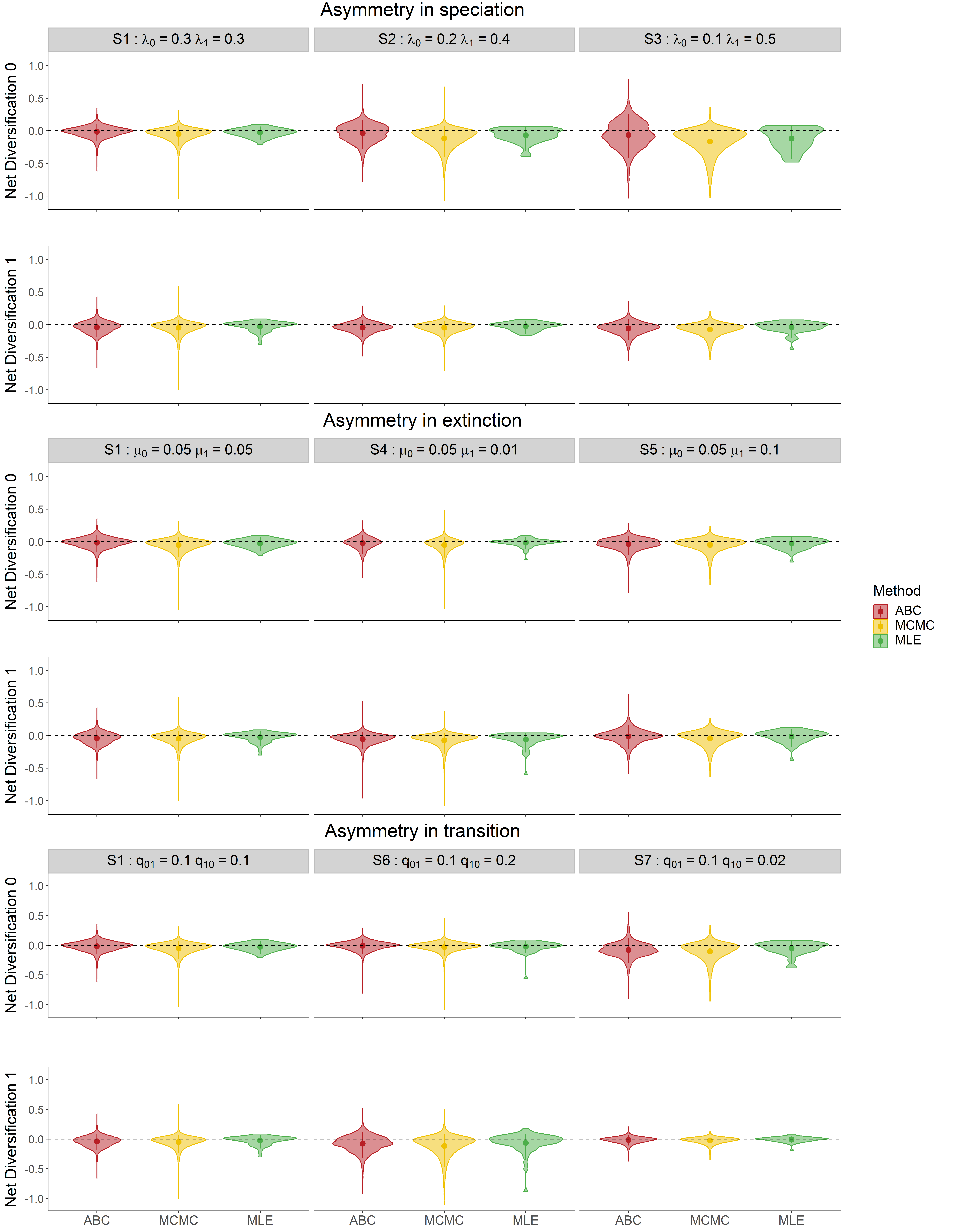

Figure S3. Parameter estimations of state-dependent **net diversification** (speciation - extinction) rates using the **ABC, MCMC and MLE** methods. The distributions shown combine the **entire posterior distribution** for each single parameter set. Plots show the residual inference error between estimated and (true) generated values. Dashed horizontal lines represent zero error to guide the eye. The colors indicate different inference methods. The ABC results are estimated using the summary statistics nLTT_total_, nLTT_0_, nLTT_1_ and *D*.

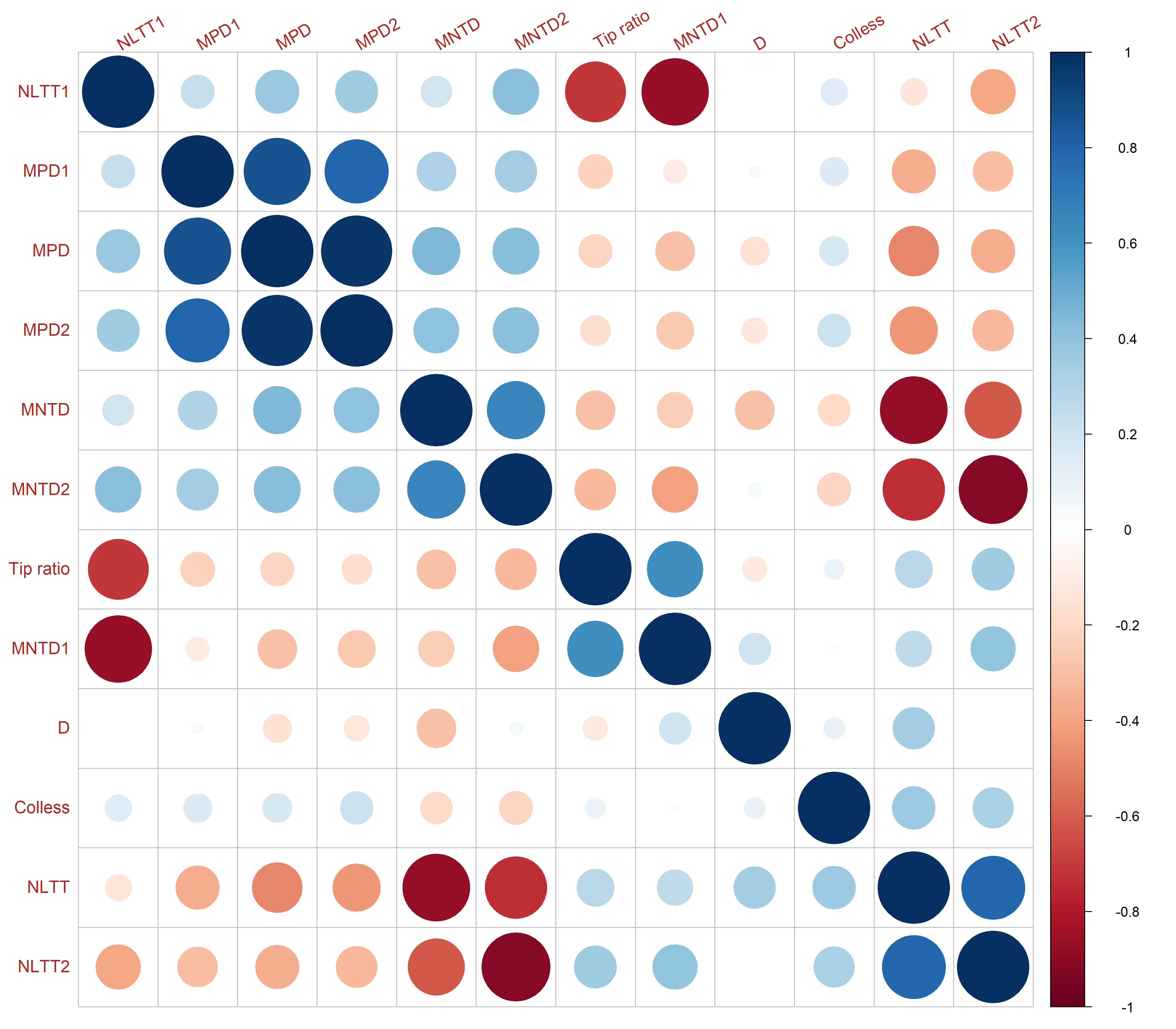

Figure S4. Correlation heatmap between summary statistics of all the observed datasets. Red indicates a negative correlation between two statistics, and blue a positive correlation. The size and color of the circle show the strength of the correlation.

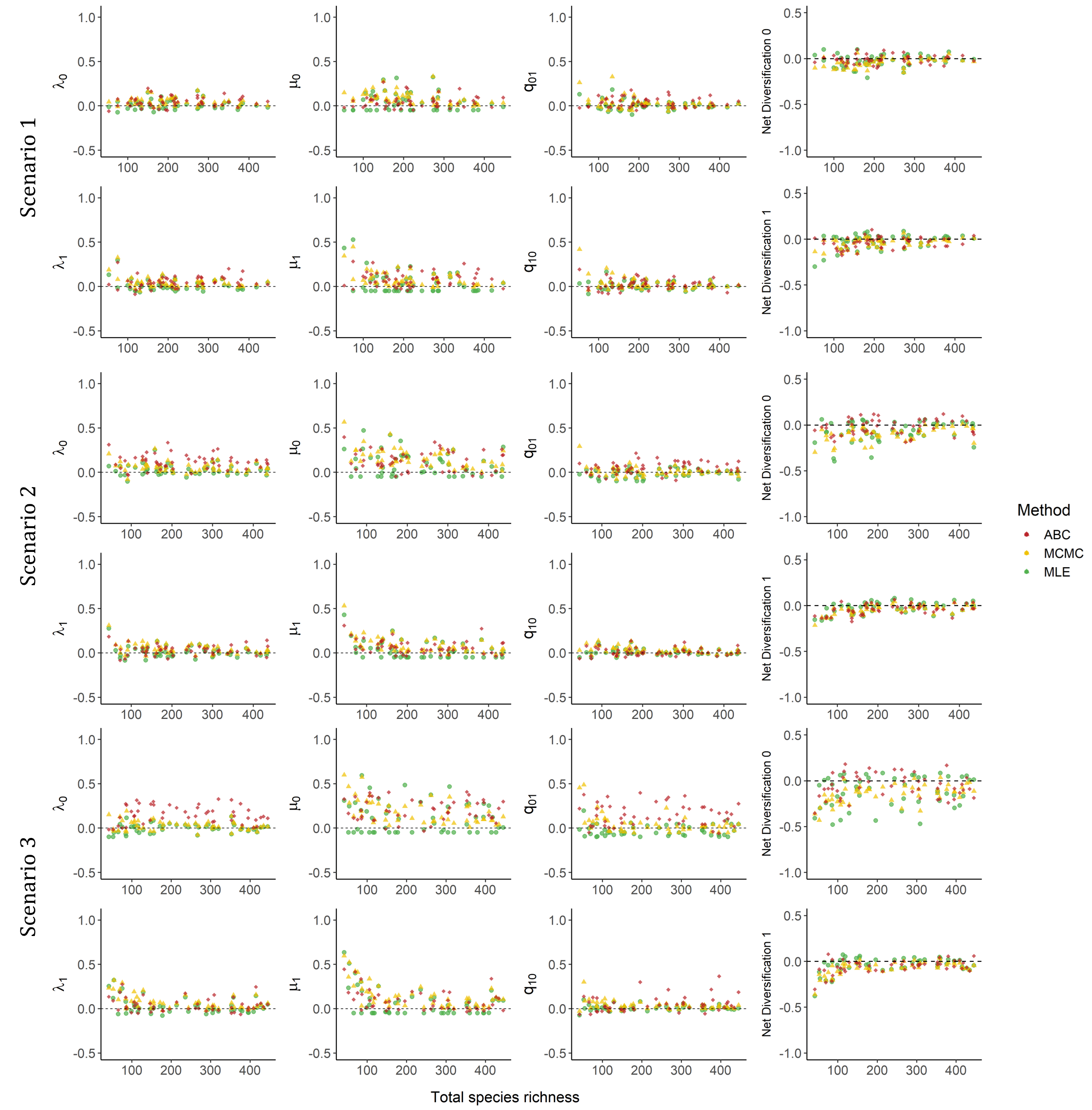

Figure S5. Parameter estimations using the **ABC, MCMC and MLE** methods, against the **total number of species** of observed trees in scenarios 1, 2 and 3. The three scenarios differ in generating rates of **speciation**, and the asymmetry level increases gradually from scenario 1 to scenario 3. The colors indicate different inference methods. Each plot has 50 points in each color, representing 50 replicates for each scenario.
